# TREM2 interacts with TDP-43 and mediates microglial neuroprotection against TDP-43-related neurodegeneration

**DOI:** 10.1101/2021.07.28.454242

**Authors:** Manling Xie, Yong U. Liu, Shunyi Zhao, Lingxin Zhang, Dale B. Bosco, Yuan-Ping Pang, Jun Zhong, Udit Sheth, Yuka A. Martens, Na Zhao, Chia-Chen Liu, Yongxian Zhuang, Liewei Wang, Dennis W. Dickson, Mark P. Mattson, Guojun Bu, Long-Jun Wu

## Abstract

Triggering receptor expressed on myeloid cell 2 (TREM2) is a surface receptor that, in the central nervous system, is exclusively expressed on microglia. TREM2 variants have been linked to increased risk for neurodegenerative diseases, but the functional effects of microglial TREM2 remain largely unknown. To this end, we investigated TAR-DNA binding protein 43 kDa (TDP-43)-related neurodegenerative disease via viral-mediated expression of human TDP-43 protein (hTDP-43) in neonatal and adult mice or inducible expression of hTDP43 with defective nuclear localization signals in transgenic mice. We found that TREM2 deficiency impaired microglia phagocytic clearance of pathological TDP-43, and enhanced neuronal damage and motor function impairments. Mass cytometry analysis revealed that hTDP-43 induced a TREM2-dependent subpopulation of microglia with high CD11c expression and higher phagocytic ability. Using mass spectrometry and surface plasmon resonance analysis, we further demonstrated an interaction between TDP-43 and TREM2, *in vitro* and *in vivo,* in hTDP-43-expressing transgenic mouse brains. We computationally identified the region within hTDP-43 that interacts with TREM2 and observed the potential interaction in ALS patient tissues. Our data reveal the novel interaction between TREM2 and TDP-43, highlighting that TDP-43 is a possible ligand for microglial TREM2 and the interaction mediates neuroprotection of microglial TREM2 in TDP-43-related neurodegeneration.

## Introduction

Microglia account for 5-10% of glial cells and serve as the principal immune cells of the central nervous system (CNS). Initially implicated as inflammatory cells, microglia are now recognized to play neuroprotective roles in neurodegenerative diseases, by sensing their environment, clearing injury stimuli via phagocytosis and preserving neuronal health ^1^. Triggering receptor expressed on myeloid cell 2 (TREM2) is exclusively expressed on microglia in the CNS and is critical for microglial proliferation, migration, and phagocytosis ^2^. Multiple lines of evidence support the critical role of TREM2 in neurodegenerative diseases. In particular, TREM2 variants are genetically linked to increased risk for Alzheimer’s disease (AD) ^3, 4^. In an AD mouse model, TREM2 was demonstrated to act as a sensor for anionic lipids associated with amyloid-β (Aβ) accumulation or degenerating neurons ^5^. In addition, recent studies have further demonstrated, both *in vivo* and *in vitro,* that TREM2 is a receptor for Aβ ^6^.

TAR-DNA binding protein 43 kDa (TDP-43) was first identified in 1995 as a DNA binding protein and plays a critical role in regulating gene expression ^7^. It is the main component of the insoluble and ubiquitinated protein aggregates found in most ALS patients ^8^. TDP-43 aggregates have also been identified in other neurodegenerative diseases, such as frontotemporal dementia (FTD) and AD ^9^. Therefore, understanding the basis of neurotoxicity related to TDP-43 aggregates might be pertinent for CNS neurodegenerative diseases in general. In epidemiologic studies, whether TREM2 variants are risk factors for ALS is still under debate. A large population based study revealed that TREM2 variant R47H increases the risk for sporadic ALS ^10^. However, other groups found that TREM2 R47H is not associated with ALS ^11^. Notably, all these studies focused on TREM2 variant R47H, while other TREM2 variants are largely understudied. Nevertheless, spatial gene expression analysis in both ALS mouse model and sporadic ALS patients implicates TREM2-mediated mechanism in disease pathogenesis ^12, 13^. In addition, the level of soluble TREM2 in cerebrospinal fluid (CSF) of ALS patients was shown to have a strong positive correlation with disease progression in late stage ^14^. However, whether or how TREM2 participates in TDP-43-related neurodegeneration remains unknown.

Towards this end, we used viral-transduction to overexpress human TDP-43 (hTDP-43) in the mouse CNS as well as a transgenic mouse model that inducibly expresses hTDP43 bearing defective nuclear localization signal (rNLS8) ^15^. We found that TREM2 mediates microglial phagocytic clearance of pathological hTDP-43 proteins and that TREM2 deficiency facilitates motor neuron degeneration. We further demonstrated that microglial TREM2 interacts with pathological hTDP-43, suggesting a potential molecular mechanism underlying the neuroprotective function for microglia in TDP-43-mediated neurodegeneration.

## Results

### TREM2 deficiency aggravates hTDP-43-induced behavioral deficits and neurodegeneration

Intracerebroventricular (ICV) injection of adeno-associated virus (AAV) into postnatal day 0 (P0) mice results in widespread transduction of neurons in the brain. We injected AAV9 into P0 neonatal mice to express hTDP-43 protein fused with a GFP tag (AAV9.CAG.hTDP43-GFP) (**Fig. 1a**). In this neonatal ICV injection model, the hTDP-43 protein was widely expressed throughout the brain and spinal cord 21 days after AAV injection, including motor cortex, hippocampus, thalamus and brainstem (**Extended Data Fig. 1a-d**). Immunostaining for NeuN, Iba1, GFAP and CNPase (markers for neurons, microglia, astrocytes and oligodendroglia, respectively) indicated that hTDP-43 was exclusively expressed in neurons (**Extended Data Fig. 1e, f**). Moreover, in line with the clinical characteristics of hTDP-43 pathology, we found hTDP-43 expression in both the nucleus and cytoplasm of neurons 5 weeks after AAV injection (**Fig. 1b, c**). In addition, pathological inclusions of phosphorylated hTDP-43 (p-hTDP-43) were also detected using p409/410 antibodies at this time point, with more severe pathology in motor cortex (**Fig. 1d, Extended Data Fig. 1g**). However, typical ubiquitinated inclusions reported in ALS patients were not detected in the neonatal ICV injection model (data not shown). We also observed significant motor deficits, as illustrated by dramatically impaired hindlimb clasping in hTDP-43 expressing mice at day 14 when compared with control mice (injected with AAV9.GFP) (**Fig. 1e**). These findings indicate that neonatal ICV injection of hTDP-43 virus can partially induce typical TDP-43 pathology and motor deficits in WT mice.

**Fig 1.**
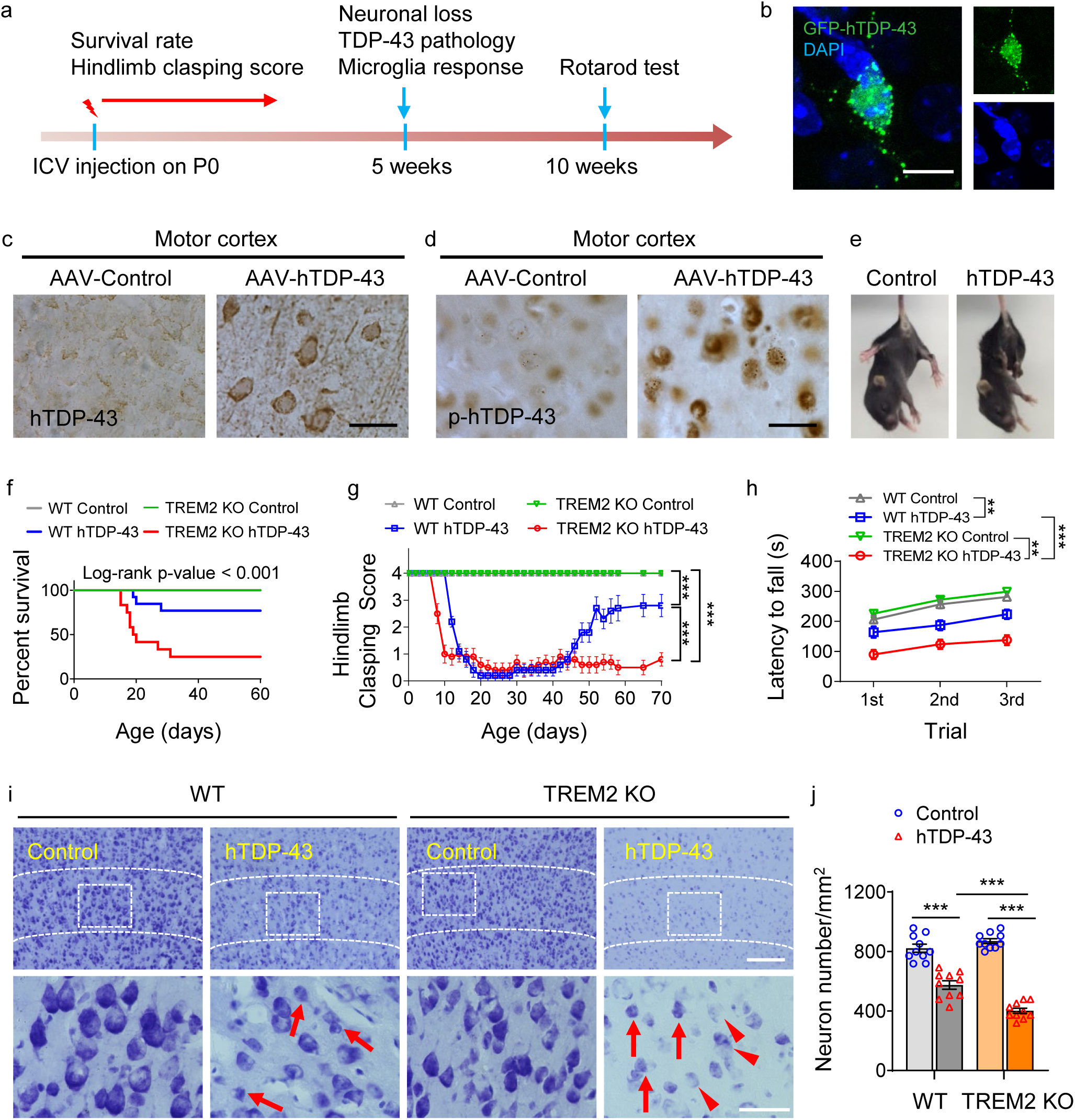
TREM2 deficiency aggravates hTDP-43-induced behavioral deficits and neurodegeneration. GFP tagged hTDP-43 protein (GFP-hTDP-43) expression was induced via intracerebroventricular injection of AAV9.CAG.hTDP43-GFP in neonatal mice (AAV9.CAG.GFP as control). **a,** Study design and timeline for the neonatal ICV injection model. **b,** Representative images of GFP-hTDP-43 expression in both nucleus and cytosol of neurons in the primary motor cortex of WT mice at 35 days post-infection (dpi); Scale bar, 10 µm. **c-d**, Representative images of hTDP-43 (**c**) or p-hTDP-43 (**d**) expression in the motor cortex at 35 dpi after AAV-control or AAV-hTDP-43 infection; Scale bar, 20 µm. **e,** Representative images of typical clasping phenotype in WT mice expressing hTDP-43 at 14 dpi. **f,** Kaplan-Meier survival curves show the percentage of mice alive at each postnatal day up to 60 dpi (n = 20 per group). **g,** Hindlimb clasping response scores collected over 70 days (n = 10 per group). **h,** Average latency to fall during rotarod analysis at 70 dpi (n=10 per group). **i,** Representative images of Nissl staining in the primary motor cortex of indicated groups at 35 dpi; Dashed lines indicate the borders of layer 4&5; Scale bar, 100 μm; High magnification images as indicated by area in dotted white box showing neuronal morphology of the cortical layer V at the bottom; Scale bar, 50 μm. Red arrows indicate neuronal shrinkage and red arrowheads indicate a significant loss of Nissl staining. **j,** Quantification of neuron numbers in the primary motor cortex of indicated groups at 35 dpi (n=10 per group). Data represented as mean ± SEM. Significance was calculated using two-way ANOVA followed by Tukey’s *post hoc* test; *n.s.*, not significant; \**P* < 0.05, \*\**P* < 0.01, *** *P* < 0.001.

To evaluate the function of TREM2 in hTDP-43-related motor deficits, we compared behavioral outcomes in TREM2 knockout (KO) and wild-type (WT) mice expressing hTDP-43. We found that mortality after hTDP-43 overexpression was significantly higher in TREM2 KO mice than that in WT mice (**Fig. 1f**). Additionally, TREM2 KO mice expressing hTDP-43 exhibited hindlimb clasping 5 days earlier than WT mice (**Fig. 1g**). Of note, TREM2 KO mice remained hunched in the ensuing observation period, while this behavioral trait began to recover in WT mice approximately 40 days post-virus injection (**Fig. 1g**). These results indicate worse hTDP-43 related motor dysfunction in TREM2 KO mice when compared with WT mice.

At 5 weeks post-virus injection, WT and TREM2 KO mice have similar hindlimb clasping response (**Fig. 1g**) and weight loss (**Extended Data Fig. 2a).** However, we found that locomotor impairment was significantly greater in TREM2 KO mice than in WT mice (reduced travel distance in open field testing, **Extended Data Fig. 2b, c**). Interestingly, travel distance by WT mice expressing hTDP-43 was slightly greater than in WT mice expressing control virus (**Extended Data Fig. 2b, c**). A similar behavioral phenotype was reported for the rTg4510 mouse model of tauopathy ^16^. Rotarod performance during the recovery phase (10 weeks post-injection) also revealed greater impairment of motor function (*i.e.*, reduction in latency to fall) in hTDP-43 expressing TREM2 KO mice than in WT mice (**Fig. 1h**). To study the association of hTDP-43-induced motor impairment with neurodegeneration, we performed Nissl staining at 5 weeks post-virus injection. Shrinkage of neuronal cell populations was detected in both WT and TREM2 KO mice expressing hTDP-43, with lower neuronal numbers in TREM2 KO mice (**Fig. 1i, j**). Similarly, NeuN immunostaining showed that NeuN loss in the primary motor cortex was greater in TREM2 KO mice than in WT mice expressing hTDP-43 **(Extended Data Fig. 2d, e**).

To determine whether motor function recovery at later time points correlated with levels of pathological hTDP-43 protein, we isolated soluble and sarkosyl-insoluble fractions from whole brains to measure the total hTDP-43 and phosphorylated hTDP-43 (p-hTDP-43) protein levels at 10 weeks post-virus injection (**Extended Data Fig. 3a**). Pathological TDP-43 aggregates resist solubilization in sarkosyl ^17^. p-TDP-43 has been reported to be one of the major components of the aggregated inclusions critical for ALS progression ^18, 19^. Western blot analysis revealed that levels of both hTDP-43 and p-hTDP-43 were much higher in TREM2 KO mice when compared to WT mice in the soluble fraction (**Extended Data Fig. 3b**). Phospho-Ser409/410 antibody also detected a ∼20 kD fragment in the soluble fractions (**Extended Data Fig. 3b**). Notably, Ser409/410 is at the extreme C-terminal of TDP-43 protein, suggesting that either the full length hTDP-43 was cleaved to generate C-terminal fragments in TREM2 KO mice, or the C-terminal fragments were less efficiently cleared by TREM2 KO mice. The sarkosyl-insoluble fraction in brains of TREM2 KO mice also contained more p-hTDP-43 (**Extended Data Fig. 3c**). Together, these results indicate that the higher level of pathological hTDP-43 in the brains of TREM2 KO mice likely contributes to their more severe, relatively irreversible motor dysfunction compared with WT mice.

### TREM2 deficiency abolishes the hTDP-43-induced CD11c^+^ microglia subpopulation

As the CNS resident immune cells, microglia undergo significant phenotypic changes following CNS injury and neurodegeneration ^20^. We therefore evaluated the effects of TREM2 deficiency on microglial phenotypes in response to hTDP-43-related neurodegeneration. We employed an integrated set of cell surface antibodies to enable mapping of microglial phenotypes via mass cytometry (CyTOF), a high dimensional single cell analysis method that can identify functional cell subpopulations ^21^. Antibodies to surface antigens CD11b, CD45, CX3CR1, CD11c, CD64, CD169, CD86, CD206, F4-80, MERTK, MHC class II, Siglec H, Ly6C, Ly6G and CCR2 as well as intracellular antigens Iba1, IL-1α, TGFβ, MCP-1, BDNF, TNFα and pNFκB were used for mass cytometry (Supplementary Table1). The microglial subpopulations and phenotypic changes were further analyzed by immunohistochemistry (**Fig. 2a**).

**Fig 2.**
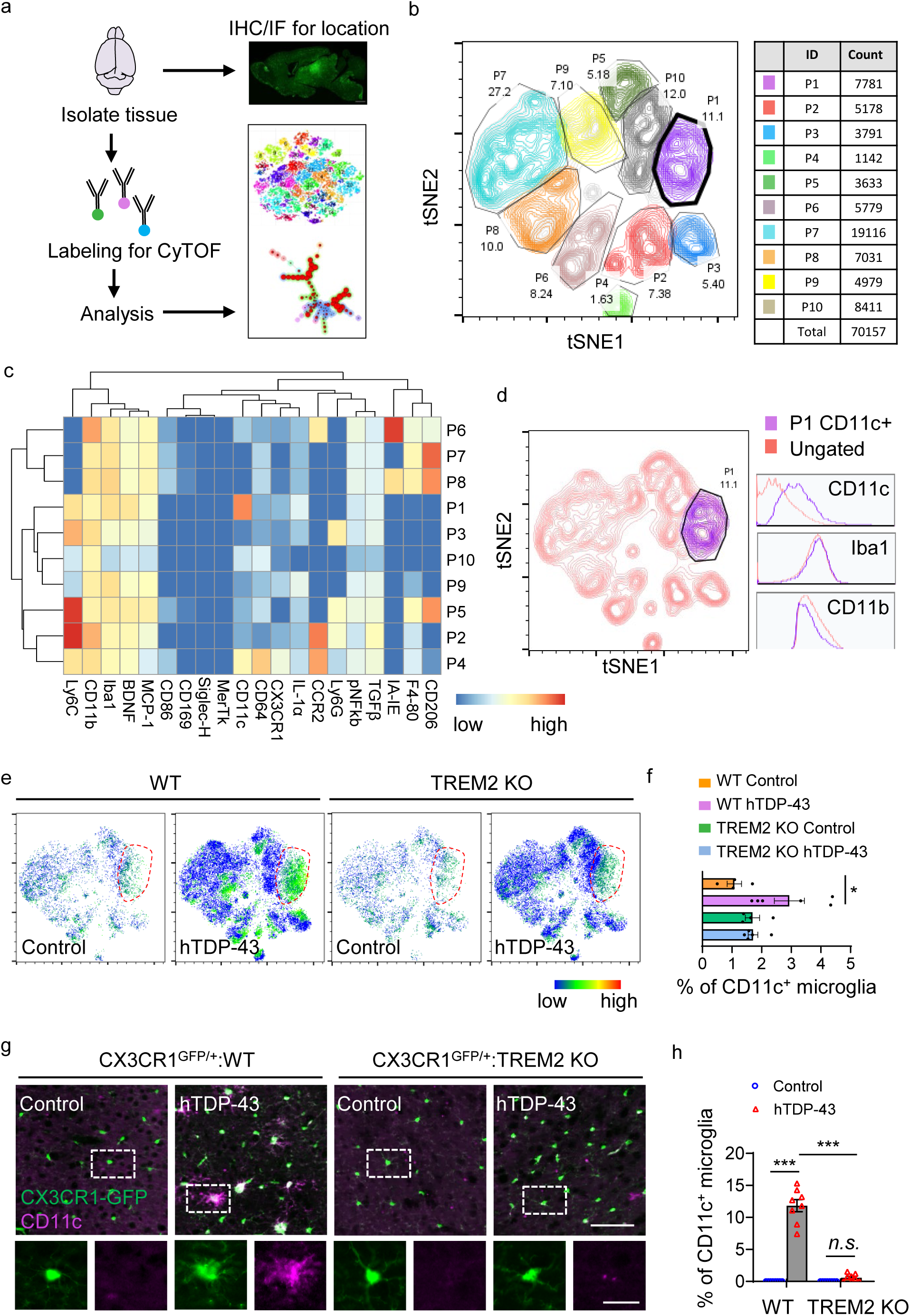
TREM2 deficiency abolishes hTDP-43-induced CD11c^+^ microglia subpopulation. hTDP-43 protein expression was induced via intracerebroventricular injection of AAV9.CAG.hTDP-43 in neonatal mice (AAV9.CAG.Empty as control). **a,** Schematic workflow for cytometry by time of flight mass spectrometry (CyTOF) and immunostaining. **b,** t-SNE map displaying 70157 CD45^med^CD11b^+^ microglia from 20 mice brain of 4 indicated groups at 35 days post-infection (dpi). Colors correspond to FlowSOM-guided clustering of microglia subpopulations (n = 4-6 per group). **c,** Heat map shows the expression levels of markers used for the microglia subpopulation analysis. Heat colors of expression levels have been scaled for each marker (blue, low expression; red, high expression). **d,** Single-cell t-SNE maps highlight CD11c^+^ microglia sub-population. The histogram panels show the expression of CD11c, Iba1 and CD11b. **e,** t-SNE map reveals a unique CD11c^+^ microglia sub-population indicated by dotted circle in WT mice expressing hTDP-43 at 35 dpi. CD11c^+^ microglia sub-population was largely abolished in hTDP-43 expressing TREM2 KO mice. The color code shows the expression level of CD11c (blue, low expression; red, high expression). **f,** Frequency analysis of CD11c^+^ microglia based on manual gating of indicated groups at 35 dpi (n = 4-6 per group). **g,** Representative images of CD11c (purple) expression by immunostaining in the primary motor cortex of indicated groups at 35 dpi; Scale bar, 100 µm. High magnification images as indicated by area in dotted white box showing the single microglia at the bottom; Scale bar, 10 µm. **h**, Average frequency of microglia expressing CD11c in the primary motor cortex of indicated groups at 35 dpi (n = 8 per group). CD11c microglia subpopulation was present in the WT mice expressing hTDP-43; however, few in TREM2 KO mice expressing hTDP-43 were observed. Data represented as mean ± SEM. Significance was calculated using two-way ANOVA followed by Tukey’s *post hoc* test; *n.s.*, not significant; \**P* < 0.05, \*\**P* < 0.01, *** *P* < 0.001.

For CyTOF analysis, we gated CD45^med^CD11b^+^ microglia from each sample for further analysis (**Extended Data Fig. 4a**). A major microglia population was characterized by CD45^mid^:CD11b^+^:CX3CR1^hi^:F4/80^+^:CD64^+^:MERTK^+^:Siglec-H^+^:CD11c^-^ signature (**Extended Data Fig. 4b**). Notably, an increased level of Iba1, CD206, CD11c, F4-80 and TGFβ and a decreased CX3CR1 level was observed in WT mice expressing hTDP-43 but not in TREM2 KO mice (**Extended Data Fig. 4c**). Frequency analysis of microglia based on manual gating revealed that numbers of microglia in the primary motor cortex were similarly increased in both WT and TREM2 KO mice expressing hTDP-43 at 5 weeks post-virus injection (**Extended Data Fig. 4d**). These results are consistent with hTDP-43-increased microglia number using immunostaining analysis in CX3CR1^GFP/+^:WT and CX3CR1^GFP/+^:TREM2 KO mice (**Extended Data Fig. 5a, b**). Transgenic CX3CR1^GFP/+^ mice expressing GFP under the control of the fractalkine promoter allow better visualization of microglial morphology ^22^. Quantitative analyses revealed that microglia differed morphologically, with TREM2 KO mice exhibiting less evidence of activation in response to hTDP-43 (*i.e.,* they had smaller soma size; **Extended Data Fig. 5c**).

We then performed visual inspection with t-SNE map (t-distributed Stochastic Neighbor Embedding) using 70157 microglia pooled from all samples for subpopulation analysis (**Fig. 2b**). Microglia were divided into 10 clusters and a detailed profile of each microglia cluster was described by phenotypic heat maps (**Fig. 2c**). Interestingly, a sub-cluster of microglia had a distinctive CD11c^high^ phenotype, which was only found in WT hTDP-43 expressing mice but not in WT control mice (**Fig. 2d-f**). Moreover, in TREM2 KO hTDP-43 expressing mice, CD11c^high^ microglial subpopulation was absent (**Fig. 2e, f**). CD11c^+^ microglia have been reported in a mouse model of AD as an activated subpopulation that clusters around amyloid plaques ^23^. We next evaluated the brains of hTDP-43 expressing mice for the distribution of CD11c^+^ microglia by immunostaining (**Fig. 2g**). Indeed, we found a highly activated microglia subpopulation expressing CD11c in the primary motor cortex of WT mice after hTDP-43 overexpression, while there were few CD11c^+^ microglia in TREM2 KO mice expressing hTDP-43 (**Fig. 2g, h**). To further explore the function of CD11c^+^ microglia, we performed co-staining for Iba1 and CD11c with GFP-hTDP-43. We observed that microglia with phagocytosed GFP-hTDP-43 (phagocytic microglia) had higher CD11c expression and accounted for 7% of total microglia (**Extended Data Fig. 5d, e**). Together, our results suggest that TREM2 deficiency abolishes the hTDP-43-induced CD11c^+^ subpopulation of microglia, which plays a critical role in phagocytosing hTDP-43.

### TREM2 deficiency locks microglia into a homeostatic status in TDP-43 neurodegeneration

To investigate how TREM2 deficiency might affect microglial response, we stereotactically injected GFP-hTDP-43 virus into the primary motor cortex of 2-month-old mice and performed a comprehensive analysis of microglial morphology and phagocytosis (**Fig. 3a and Extended Data Fig. 6a**). At day 14 in this adult local injection model, hTDP-43 inclusions were found predominantly in neuronal nuclei and diffusely in cytoplasm (**Extended Data Fig. 6b**). hTDP-43 expression in WT and TREM2 KO groups did not differ significantly at day 14, as revealed by the density analysis of GFP-hTDP-43 particles (**Extended Data Fig. 6c, d**), indicating similar viral transduction efficiency. This is further confirmed by immunoblotting against GFP or TDP-43 (**Extended Data Fig. 6e**). As expected, microglia accumulation was abundant in areas of hTDP-43 expression at 14 days post-virus injection of both WT and TREM2 KO mice (**Extended Data Fig. 6f, g**). In addition, WT microglia exhibited a canonical reactive phenotype, less ramified with enlarged soma and shorter processes in response to hTDP-43 expression (**Fig. 3b-e**). However, this hTDP-43-induced reactive phenotype was significantly attenuated in TREM2 KO mice (**Fig. 3b-e**). These data indicate that TREM2 deficiency reduces microglial activation following TDP-43 neurodegeneration.

**Fig 3.**
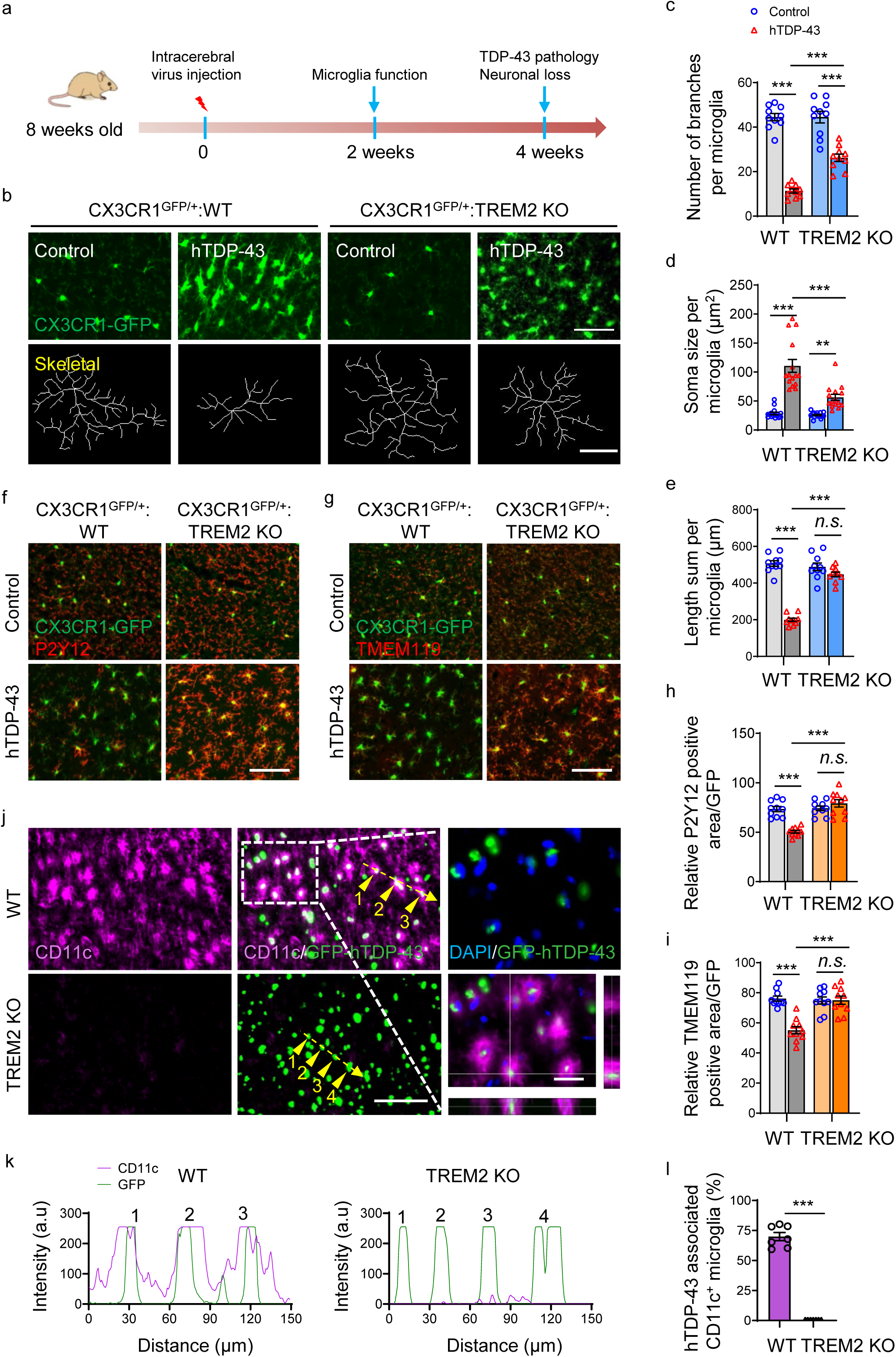
TREM2 deficiency locks microglia into a homeostatic state in TDP-43-induced neurodegeneration. In adult local injection model, hTDP-43 or GFP-hTDP-43 was expressed in the primary motor cortex of 2-month-old mice via stereotactic intracerebral injection of AAV9.CAG.hTDP-43 or AAV9.CAG.hTDP43-GFP (AAV9.CAG.Empty or AAV9.CAG.GFP as control). **a,** Study design and timeline for local hTDP-43 expression model. **b,** Representative images (upper panels; Scale bar, 100 µm) and transformed skeletal images (lower panels; Scale bar, 10 µm) of GFP-expressing microglia in the primary motor cortex of indicated groups at 14 dpi. **c-e,** Quantification of GFP-expressing microglia number of branches (**c**, n=10 per group), soma size (**d**, n=15 per group) and process length (**e**, n=10 per group) in the primary motor cortex of indicated groups at 14 dpi. WT microglia displayed a reactive phenotype as reduced branches, increased soma size and shorter processes in response to hTDP-43 expression. However, this hTDP-43-induced reactive phenotype was significantly attenuated in TREM2 KO mice. **f,** Representative images of P2Y12 expression (red) in the primary motor cortex of indicated groups at 7 dpi; Scale bar, 100 µm. **g,** Representative images of TMEM119 expression (red) in the primary motor cortex of indicated groups at 7 dpi; Scale bar, 100 µm. **h, i,** Quantification of relative P2Y12 and TMEM119 positive area in the primary motor cortex of indicated groups at 7 dpi (n=10 per group). **j,** Representative images of CD11c (purple), hTDP-43 (green), and DAPI staining (blue) in the primary motor cortex of indicated groups at 28 dpi. Scale bar, 100 µm. High magnification images as indicated by area in dotted white box showing on the right; Scale bar, 10 µm. **k,** Analysis of co-localization of CD11c (purple curves) with GFP-hTDP-43 (green curves) in the primary motor cortex of indicated groups at 28 dpi. Fluorescence intensity profiles of CD11c and GFP-hTDP-43 show the distribution of fluorescence across the yellow dotted arrows in **(j)**. **l,** Average frequency of hTDP-43 associated CD11c^+^ microglia in the primary motor cortex of indicated groups at 28 dpi (n = 7 per group). Results indicate that CD11c^+^ microglia subpopulation was present in WT mice upon hTDP-43 expression. However, CD11c^+^ microglia were not observed in TREM2 KO mice expressing GFP-hTDP-43. Data are represented as mean ± SEM. Significances were calculated using Two-way ANOVA, Tukey’s post-hoc analysis (**c**-**e**, **h** and **i**) and student t test (**l**); *n.s.*, not significant; \**P* < 0.05, \*\**P* < 0.01, *** *P* < 0.001.

We next examined two microglia homeostatic markers, P2Y12 and TMEM119 ^24^, in response to hTDP-43 expression. P2Y12 and TMEM119 were largely confined to microglial processes under basal conditions in both WT and TREM2 KO mice (**Fig. 3f, g**). We found that hTDP-43 induced remarkable downregulation of P2Y12 and TMEM119 in WT microglia (**Fig. 3f-i**). However, both P2Y12 and TMEM119 expression were unchanged from baseline in TREM2 KO microglia in response to hTDP-43 (**Fig. 3f-i**). These results suggest that TREM2 deficiency attenuates microglial activation and locks microglia into a homeostatic status in TDP-43 neurodegeneration.

We further characterized CD11c^+^ microglia subpopulation in this adult local injection model. Consistent with the neonatal ICV injection model, we found a dramatic upregulation of CD11c in microglia from WT mice in response to local hTDP-43 expression. In addition, CD11c^+^ microglia subpopulation was not observed in TREM2 KO mice expressing GFP-hTDP-43 (**Fig. 3j**). Notably, the percentage of CD11c+ microglia is much higher in adult local injection model compare with neonatal ICV injection because of more concentrated pathology in the adult local injection model (while there is relatively diffused TDP-43 pathology in the neonatal ICV model). More importantly, we found that CD11c^+^ microglia are mostly associated with GFP-hTDP-43 in WT mice but not in TREM2 KO mice by co-localization analysis (**Fig. 3k, l**). These results demonstrate that TREM2-dependent CD11c^+^ microglia phagocytose pathological hTDP-43 proteins, which manifests the critical function of TREM2 in microglia reactive phenotype towards TDP-43 neurodegeneration.

### TREM2 deficiency reduces clearance of pathological hTDP-43 by microglia

A recent study suggested that microglia actively participate in clearing hTDP-43 in a transgenic TDP-43 mouse model ^25^. Additionally, TREM2 is important for phagocytosis of cell debris from apoptotic neurons and amyloid plaques by microglia ^26, 27^. We therefore tested whether TREM2 deficiency affects microglia-mediated hTDP-43 clearance. To this end, we first performed immunostaining for CD68, a lysosomal marker upregulated in phagocytic cells, at 2 weeks following hTDP-43 expression in the primary motor cortex. Our results showed that hTDP-43 induced a dramatic increase of CD68 expression in WT but not in TREM2 KO mice (**Fig. 4a**). The analysis of CD68 and CX3CR1-GFP co-localization further confirmed the upregulation of CD68 in the individual microglia was reduced in TREM2 KO mice (**Fig. 4b**). In addition, these CD68^+^ microglia have high CD11c expression and are co-localized with GFP-hTDP-43 (**Fig. 4c**). Next, we directly investigated whether hTDP-43 was phagocytosed by microglia. We tagged hTDP-43 with GFP and stained microglial with Iba1. At 4 weeks following hTDP-43 expression, triple-immunostaining of DAPI, Iba1 and GFP-hTDP-43 showed that hTDP-34 were in the cytoplasm but not nucleus of Iba1^+^ microglia (**Fig. 4d**). A large number of GFP-hTDP-43 were present in Iba1^+^ microglia in WT mice, but were rarely observed inside Iba1^+^ microglia in TREM2 KO mice (**Fig. 4d, e, Extended Data Fig. 6h**). Co-localization analysis further confirmed that GFP-hTDP-43 existed in soma of WT microglia, but not TREM2 KO microglia (**Fig. 4f**). Consistent with the neonatal ICV injection model, we found that phagocytic microglia (colocalized with hTDP-43) had a higher CD11c expression compared with non-phagocytic microglia (**Fig. 4g, h**). These data further demonstrate that TREM2 is required for hTDP-43-induced CD11c+ subpopulation of microglia which has strong phagocytic functions.

**Fig 4.**
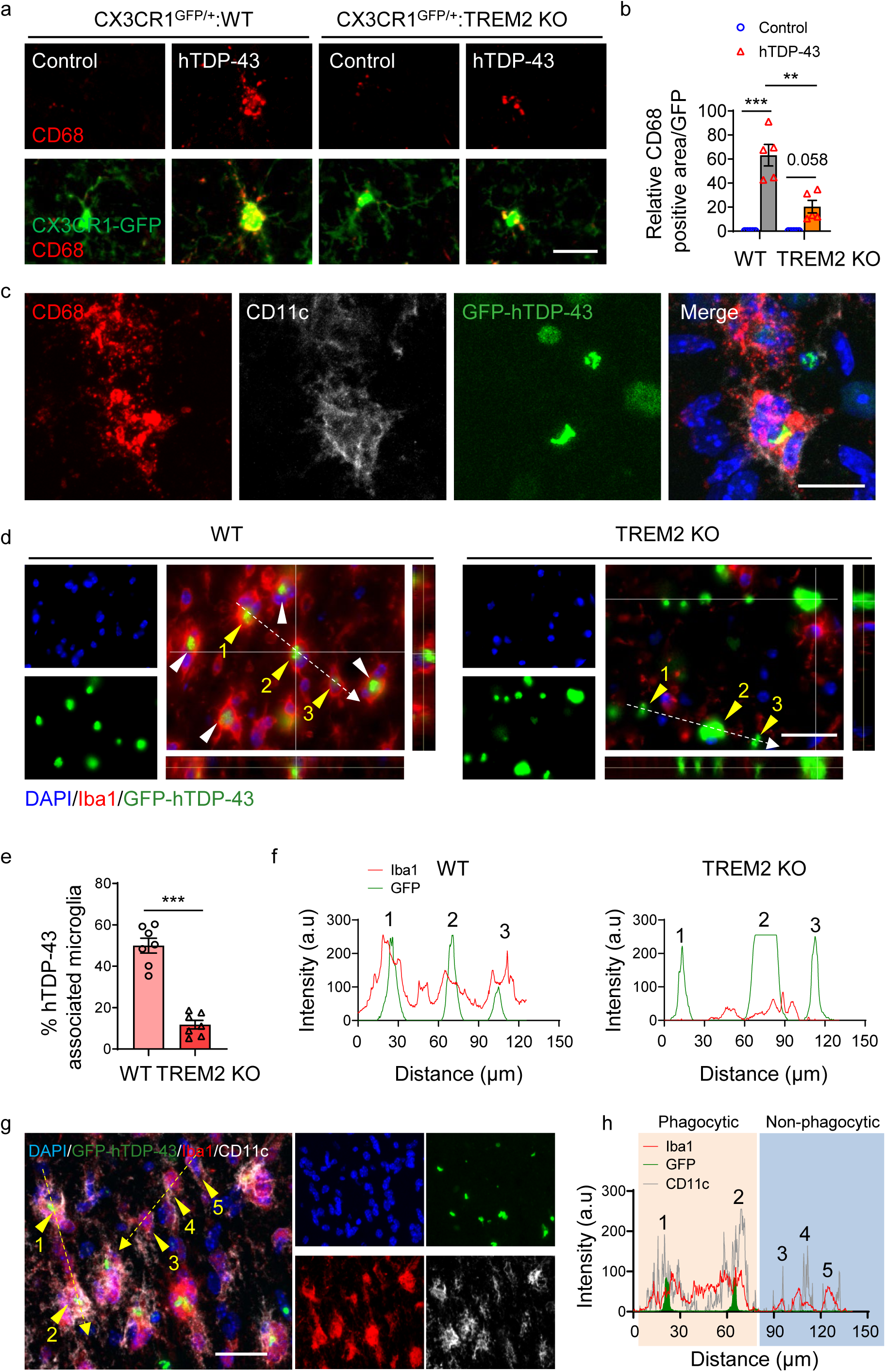
TREM2 deficiency impairs microglial phagocytosis of pathological hTDP-43 protein. hTDP-43 or GFP-hTDP-43 was expressed in the primary motor cortex of 2-month-old mice via stereotactic intracerebral injection of AAV9.CAG.hTDP-43 or AAV9.CAG.hTDP43-GFP (AAV9.CAG.Empty or AAV9.CAG.GFP as control). **a,** Representative images of CD68 expression (red) in microglia (CX3CR1-GFP, green) in the primary motor cortex of indicated groups at 14 dpi; Scale bar, 100 µm. **b,** Quantification of relative CD68 positive area in the primary motor cortex of indicated groups at 14 dpi (n=5 per group). Results show hTDP-43 induced increase of CD68 expression in WT but not in TREM2 KO mice. **c,** Representative images of CD68 (red) and CD11c (white) expression in microglia phagocytosing GFP-hTDP-43 (green) in the primary motor cortex of WT mice at 14 dpi; Scale bar, 10 µm. **d,** Representative images of microglia (Iba1, red) phagocytosis of GFP-hTDP-43 (green) in the primary motor cortex of WT but not TREM2 KO mice at 28 dpi, as indicated by the arrowheads; Scale bar, 20 µm. **e,** Quantification of hTDP-43 associated microglia (Iba1) in the primary motor cortex of WT and TREM2 KO mice at 28 dpi (n = 7 per group). Result indicates decreased hTDP-43 associated microglia in TREM2 KO mice. **f,** Analysis of co-localization of Iba1 (red curves) and GFP-hTDP-43 (green curves) in the primary motor cortex of WT and TREM2 KO mice at 28 dpi. Fluorescence intensity profiles of Iba1 and GFP-hTDP-43 show the distribution of fluorescence across the white dotted arrows in **(d)**. **g,** Representative images of co-localization of CD11c (white) with Iba1 (red) in microglia phagocytosing GFP-hTDP-43 (green) in the primary motor cortex of WT mice at 28 dpi; Scale bar, 10 µm. **h,** Analysis of co-localization of Iba1 (red curves), CD11c (grey curves) and GFP-hTDP-43 (green curves) in the primary motor cortex of WT mice at 28 dpi. Fluorescence intensity profiles of Iba1, CD11c, and GFP-hTDP-43 show the distribution of fluorescence across the yellow dotted arrows in **(g)**. Data are represented as mean ± SEM. Significances were calculated using Two-way ANOVA, Tukey’s post-hoc analysis (**b**) and student t test (**e**); *n.s.*, not significant; \**P* < 0.05, \*\**P* < 0.01, *** *P* < 0.001.

### TREM2 deficiency leads to greater brain tissue accumulation of hTDP-43 protein

To characterize the dynamic changes of hTDP-43 pathology resulting from microglial phagocytic clearance, we examined total and p-TDP-43 level at 4 weeks following hTDP-43 expression. Our immunostaining results showed that p-TDP-43 was completely co-localized with GFP-hTDP-43 clusters in both WT and TREM2 KO mice (**Fig. 5a, b**).

**Fig 5.**
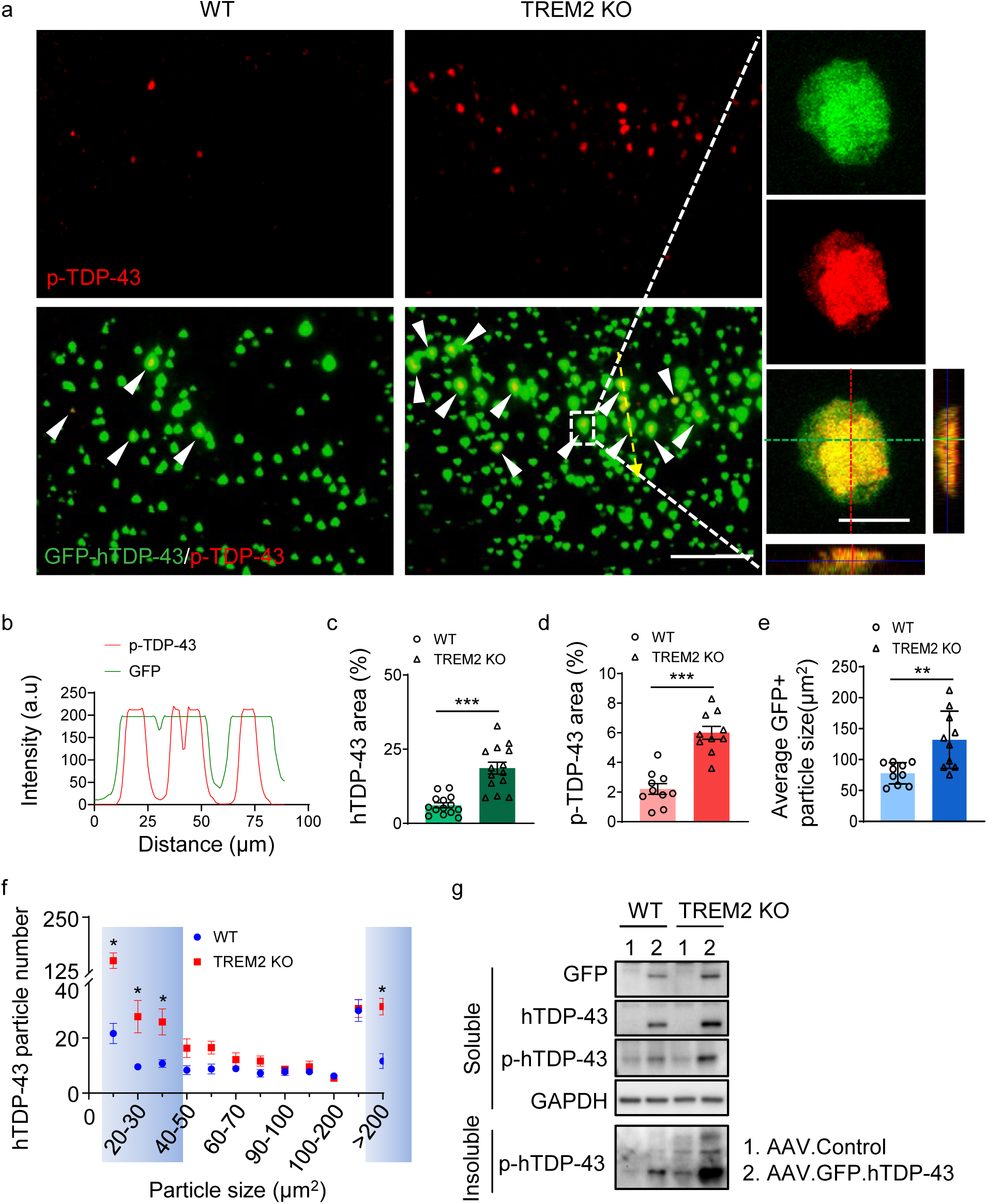
TREM2 deficiency leads to increased accumulation of pathological hTDP-43 protein. a, Representative images of co-localization of p-TDP-43 (red) with GFP-hTDP-43 in the primary motor cortex of WT and TREM2 KO mice at 28 dpi, as indicated by white arrowheads; Scale bar, 100 µm. High magnification images as indicated by area in dotted white box showing on the right; Scale bar, 10 µm. **b,** Analysis of co-localization of p-TDP-43 (red curves) and GFP-hTDP-43 (green curves) in the primary motor cortex of WT and TREM2 KO mice at 28 dpi. Fluorescence intensity profiles of p-TDP-43 and GFP-hTDP-43 show the distribution of fluorescence across the yellow dotted arrow in **(a)**. **c-e,** Quantification of hTDP-43 area (**c**, n=14 per group), p-TDP-43 area (**d,** n=10 per group) and GFP^+^ particle size (**e,** n=10 per group) in the primary motor cortex of WT and TREM2 KO mice at 28 dpi. Results indicate higher hTDP-43, p-TDP-43 and GFP^+^ in TREM2 KO mice. **f,** hTDP-43 particle size distribution in the primary motor cortex of WT and TREM2 KO mice at 28 dpi. Particles with size below 40 µm^2^ or over 200 µm^2^ are highlighted by blue (n=9 per group). **g,** GFP-hTDP-43 and p-hTDP-43 (Ser403/404) immunoblots of the soluble fractions and Sarkosyl-insoluble fraction from primary motor cortex of WT and TREM2 KO mice at 28 dpi. Western blots were independently repeated four times (n=4 per group). GAPDH was used as loading control. TREM2 KO mice show high level of GFP-hTDP-43 and p-hTDP-43 as compared to WT mice. Data represented as mean ± SEM. Significance was calculated using student t test; *n.s.*, not significant; \**P* < 0.05, \*\**P* < 0.01.

However, TREM2 KO mice had significantly more hTDP-43 and p-TDP-43 accumulation than WT mice (**Fig. 5c, d**). In addition, compared with WT mice, average size of GFP-hTDP-43 clusters was much larger in TREM2 KO mice (**Fig. 5e**). Particle size distribution analysis revealed that hTDP-43 clusters with size below 40 µm^2^ or over 200 µm^2^ were much higher in TREM2 KO than in WT mice (**Fig. 5f**).

Using Western blot against GFP or TDP-43, we consistently found the higher level of hTDP-43 in TREM2 KO mice than in WT mice in radioimmunoprecipitation assay buffer (RIPA)-soluble fractions (**Fig. 5g**). Immunoblotting against p-hTDP-43 revealed a ∼27 kD fragment in the soluble fraction, the level of which was significantly higher in TREM2 KO mice following hTDP-43 expression. Interestingly, the phosphorylated full length hTDP-43 was not detected in the adult local injection model, suggesting that the cleaved products are more prone to posttranslational modification (**Fig. 5g**). To test whether the large GFP-hTDP-43 particles with high p-TDP-43 content were aggregated, we then isolated sarkosyl-insoluble fractions from brain tissues and performed Western blot analysis. Indeed, p-hTDP-43 was significantly more abundant in the sarkosyl-insoluble fraction of TREM2 KO brains than in WT brains following hTDP-43 expression (**Fig. 5g**). Consistent with the observations from the soluble fractions, cleaved p-hTDP-43 fragments were the major component of the hyperphosphorylated hTDP-43 (**Fig. 5g**). Together, these results indicate that microglia TREM2 is critical for microglial phagocytosis and clearance of TDP-43. In TREM2 KO mice, the uncleared hTDP-43 clusters are prone to be cleaved and then form hyperphosphorylated aggregates.

We next assessed neuronal loss in the motor cortex via NeuN staining 4 weeks after hTDP-43 expression. Consistent with the neonatal ICV injection model, we found that hTDP-43 expression lead to reduced NeuN staining, which was further aggravated in TREM2 KO mice (**Extended Data Fig. 7a, b**). By co-staining with Iba1, we found that a large number of microglia in WT mice were located close to presumably damaged neurons, following hTDP-43 expression (**Extended Data Fig. 7c, d**). However, the proximity of microglia to neurons was unremarkable in TREM2 KO mice. In addition, we found that those neuron-associated microglia had high CD11c expression (**Extended Data Fig. 7e**), suggesting a phagocytic function. Thus, these results collectively demonstrate that TREM2 deficiency leads to greater brain tissue accumulation of hTDP-43 protein and aggravates TDP-43-induced neurodegeneration.

### TDP-43 interacts with TREM2 *in vitro* and *in vivo*

The TDP-43 protein in ALS patient blood and CSF is likely released from cells via extracellular vesicles ^28, 29^. We hypothesize that hTDP-43 released into the CNS parenchyma from degenerating neurons can be detected by microglia through TREM2. To test this hypothesis, we used human iPSC-derived neurons infected with GFP-hTDP-43 virus and evaluated hTDP-43 release. By day 21 after virus infection, hTDP-43 was robustly expressed throughout the cytoplasm and neuronal damage was extensive (**Extended Data Fig. 8a**). We collected the culture medium and performed Western blot to examine whether hTDP-43 was released extracellularly. Indeed, extracellular hTDP-43 was detectable in the culture medium directly or by capture via bead-immobilized GFP antibody (**Extended Data Fig. 8b**). Interestingly, by using antibody recognizing N-terminal hTDP-43, we observed cleaved products at around ∼25 kD in the culture medium input, suggesting a mix of full-length and N-terminal fragments of hTDP-43 were released into the extracellular space (**Extended Data Fig. 8b**).

Next, to test whether TREM2 can interact with hTDP-43, we performed the co-immunoprecipitation (co-IP) experiments to examine their potential interaction. To this end, we transfected human embryo kidney (HEK) 293T cells with myc-tagged hTREM2 (myc-hTREM2) or control plasmids. hTREM2 was immunoprecipitated using an hTREM2 antibody. Immunoblot against hTDP-43 revealed the co-IP of endogenous hTDP-43 from HEK cells (**Fig. 6a**). We then performed liquid chromatography with tandem mass spectrometry (LC-MS/MS) to characterize the interaction between TREM2 and TDP-43. Specifically, annotated MS/MS spectra of V^124^LVEVLADPLDHR^136^ from hTREM2 and T^103^SDLIVLGLPWK^114^ from hTDP-43 were obtained from anti-hTREM2 immunoprecipitates (**Fig. 6b**). The two peptides are a fragment of the extracellular ligand-binding domain (ECD) of hTREM2 and a fragment of the RNA recognition motif 1 (RRM1) of hTDP-43, respectively. To further explore which domain(s) of hTDP-43 interacted with TREM2, we co-transfected HEK cells with myc-hTREM2 and GFP-hTDP-43 C terminal fragment (residues 216-414) or GFP control plasmids (**Extended Data Fig. 8c**). GFP or GFP-tagged proteins were captured using bead-immobilized GFP antibody. Immunoblot using anti-Myc revealed the co-IP of GFP-hTDP-43 C terminal fragment with myc-hTREM2, indicating the interaction of hTREM2 with the hTDP-43 C-terminal fragment (**Extended Data Fig. 8d**). In addition, co-IP using bead-immobilized myc antibody obtained similar results (**Extended Data Fig. 8e**). To further distinguish hTREM2 interacting proteins from contaminating proteins non-specifically bound to the matrix, we employed a stable isotope labeling with amino acids in cell culture (SILAC)-based quantitative proteomics approach to study the hTREM2 interactome in HEK293T cells ^30, 31^ (**Extended Data Fig. 9a**). As expected, hTREM2 was completely labeled with heavy amino acid as evidenced by the lack of light version of peptides (**Extended Data Fig. 9b and c**). The analysis of mass spectra revealed 157 potential hTREM2 interacting proteins including hTDP-43, the MS spectra of which also showed the lack of light version of peptides like hTREM2. A list of the 10 proteins with highest intensity was provided in **Extended Data Fig. 9d**. Together, these results suggest that hTDP-43 is able to interact with TREM2 *in vitro*.

**Fig 6.**
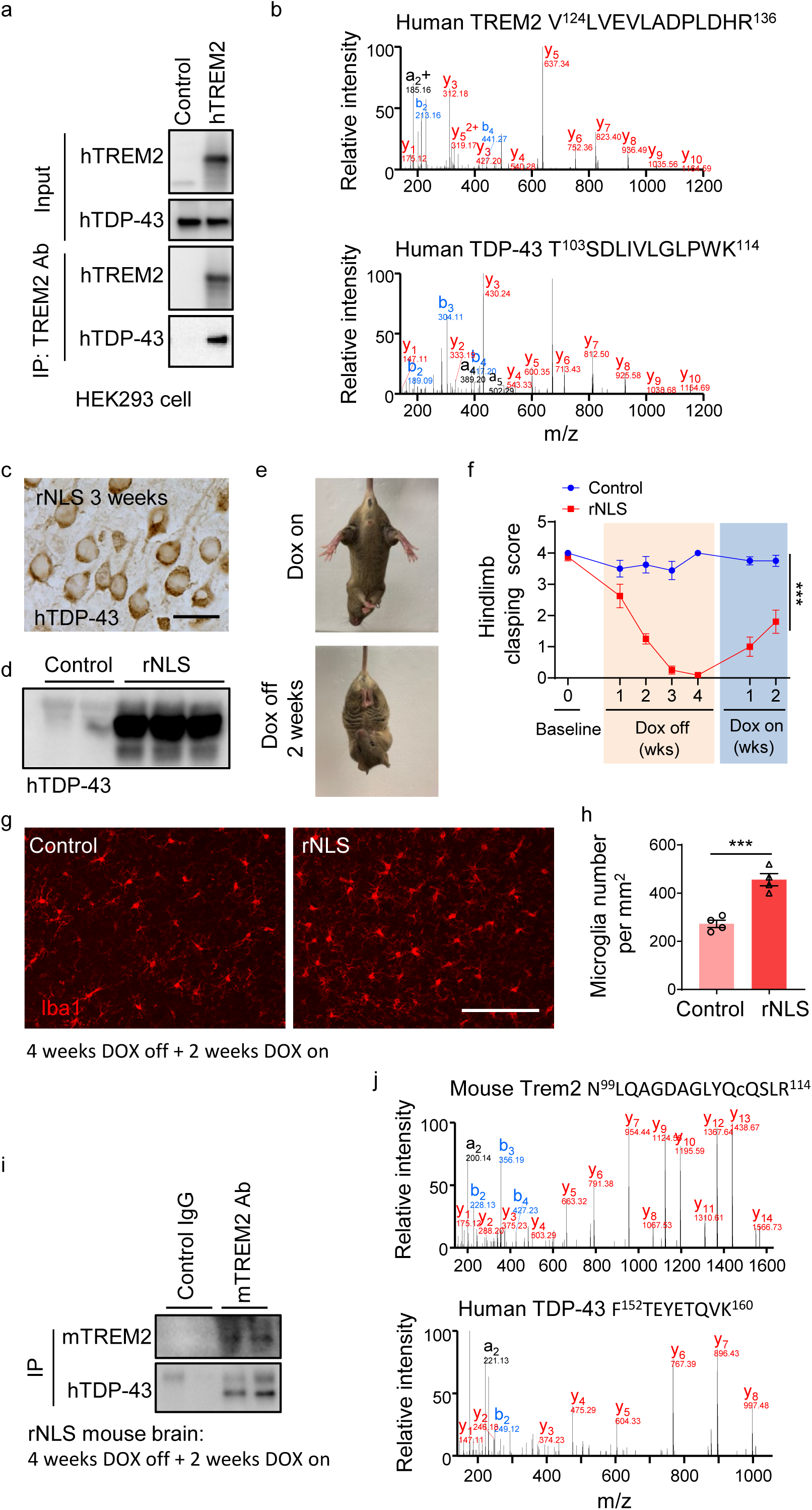
hTDP-43 interacts with TREM2 *in vitro* and *in vivo* in mouse brain. a, hTDP-43 was co-immunoprecipitated with myc-hTREM2 in HEK 293 cells overexpressing hTREM2. Experiments were independently repeated three times. **b,** Annotated MS/MS spectra of V^124^LVEVLADPLDHR^136^ from hTREM2 and T^103^SDLIVLGLPWK^114^ from hTDP-43 identified in anti-hTREM2 immunoprecipitates from HEK 293 cells. **c**, Representative images of hTDP-43 expression in cytosol in the primary motor cortex of rNLS mice at 3 weeks off DOX diet; Scale bar, 20 µm. **d,** hTDP-43 immunoblots of primary motor cortex of rNLS mice at 3 weeks off DOX diet. **e,** rNLS mice showed clasping phenotype at 2 weeks off DOX diet. **f,** Hindlimb clasping response scores over 4 weeks off DOX diet followed by 2 weeks DOX on (n = 12 per group). **g,** Representative images of microglia (Iba1, red) in the primary motor cortex of indicated groups at 4 weeks off DOX diet followed with 2 weeks DOX on. **h,** Quantification of microglial density (n=10 per group) in the primary motor cortex of indicated groups at 4 weeks off DOX diet followed by 2 weeks DOX on. Results indicate increased microglia number in rNLS mice. **i,** Endogenous mTREM2 was co-immunoprecipitated with hTDP-43 in the motor cortex of rNLS mice using mTREM2 antibody at 4 weeks off DOX diet followed with 2 weeks DOX on. Experiments were independently repeated three times. **j,** Annotated MS/MS spectra of N^99^LQAGDAGLYQcQSLR^114^ (c is carbaminomethylated cysteine) from mouse TREM2 and F^152^TEYETQVK^160^ from hTDP-43 in anti-mTREM2 immunoprecipitates from rNLS mouse cortex tissue at 4 weeks off DOX diet followed by 2 weeks DOX on. Data represented as mean ± SEM. Significances were calculated using Two-way ANOVA, Tukey’s post-hoc analysis (**f**) and student t test (**h**); *n.s.*, not significant; \**P* < 0.05, \*\**P* < 0.01, *** *P* < 0.001.

To evaluate the possible interaction between TREM2 and TDP-43 *in vivo*, we used a new inducible mouse model of ALS-rNLS8, which expresses mutant hTDP43 (hTDP43-ΔNLS) in neurons in a doxycycline (DOX)-dependent manner ^15^. We confirmed that mutant hTDP-43 accumulated in cytoplasm after expression (**Fig. 6c. d**), as the hTDP43-ΔNLS lacks nuclear localization sequence (NLS). Consistent with previous studies ^15^, we observed typical clasping phenotype when hTDP43-ΔNLS was induced during removal of DOX diet, with a rapid functional recovery as early as 2 weeks back on DOX (**Fig. 6e, f**). In addition, we found dramatic microglia activation during the recovery period (**Fig. 6g, h**). We then performed immunoprecipitation using a biotinylated mTREM2 antibody in rNLS8 mouse brain lysates. Immunoblot against mTREM2 and hTDP-43 revealed the co-IP of endogenous mTREM2 with hTDP-43 (**Fig. 6i**). We then analyzed the anti-mTREM2 immunoprecipitates from rNLS mouse cortex tissue by LC-MS/MS. Similarly, we were able to detect both mTREM2 and hTDP-43 in the mTREM2 immunoprecipitates and the MS/MS spectra of a representative peptide N99LQAGDAGLYQCQSLR114 from mTREM2 and a representative peptide F152TEYETQVK160 from hTDP-43 were shown in **Fig. 6j**, suggesting mTREM2 is physically associated with hTDP-43.

Next, we directly investigated whether TREM2 interacts with hTDP-43 *in situ*. To this end, we performed co-staining of Iba1 with TREM2 at 2 weeks following GFP-hTDP-43 expression in the adult primary motor cortex. No specific signals were detected in TREM2 KO mice, demonstrating the antibody specificity ^32^. In WT mice with hTDP-43 overexpression, we found that TREM2 is co-localized with GFP-hTDP-43, suggesting the potential interaction of TREM2 with TDP-43 *in situ* (**Extended Data Fig. 10a**).

The next plausible question was whether there are potential interactions between TREM2 and TDP-43 in human ALS patients. To this end, we obtained frozen autopsied specimens of ALS patients from Mayo Clinic Brain Bank for Neurodegenerative Disorders and examined the interaction. Basic characteristics of human tissue donors are listed in supplementary Table 2. First, in addition to well-known increased expression of TDP-43 in ALS patients ^8^, we found that TREM2 expression in both cortex and spinal cord from ALS patients was greater than those from non-neurological controls (**Fig. 7a, b**). More importantly, an endogenous hTDP-43/TREM2 complex was co-immunoprecipitated from ALS patient spinal cord tissue lysates (**Fig. 7c**). Taken together, our results indicate the interactions between TREM2 and hTDP-43 *in vivo* from both mouse and human tissues.

**Fig 7.**
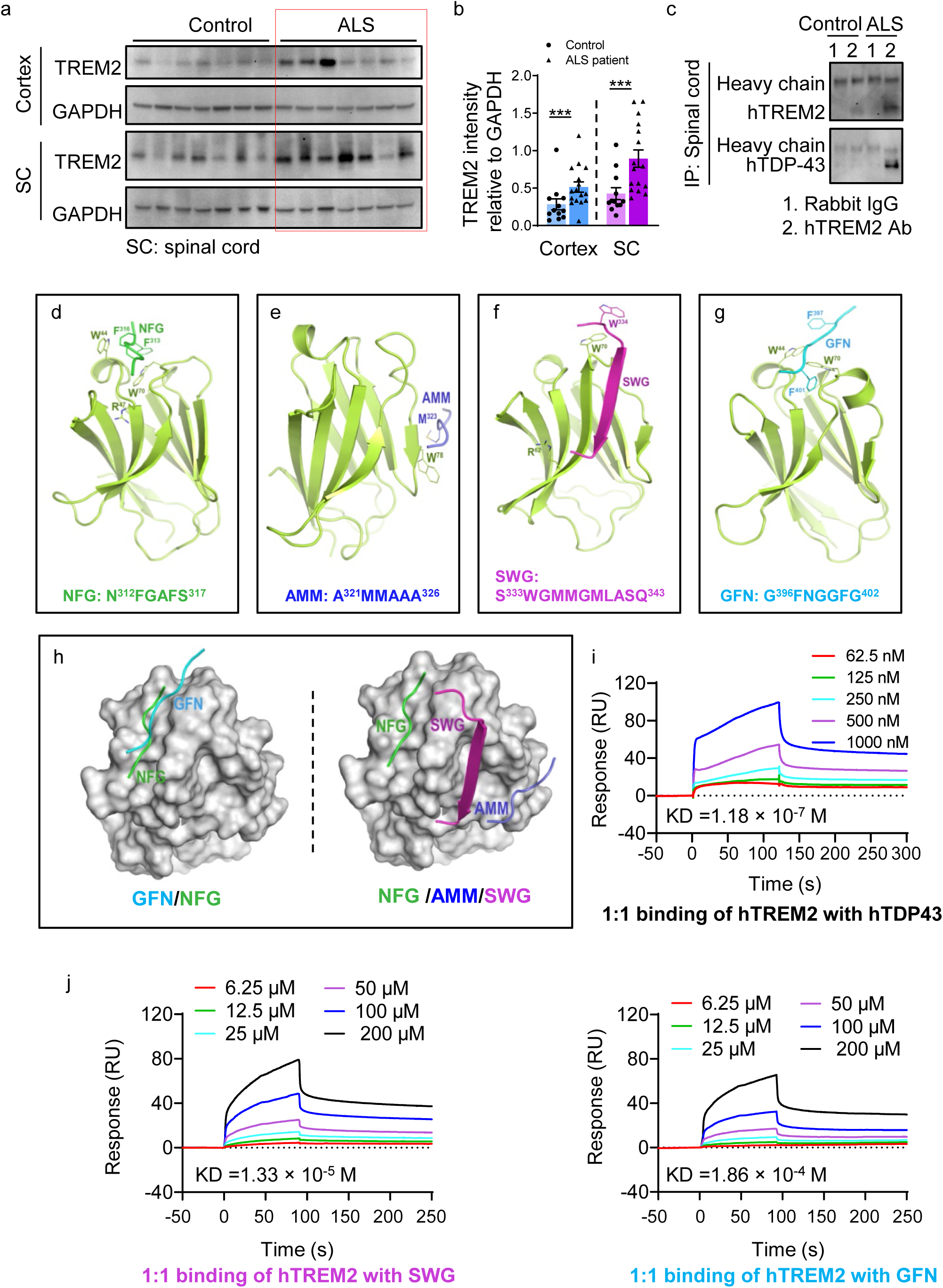
hTDP-43 interacts with TREM2 in human tissues and *in silico*. a, TREM2 immunoblots of frozen autopsied spinal cord specimens of ALS patients and age matched controls. **b,** Quantification of TREM2 level relative to GAPDH (n=12 for control, n=16 for ALS patients). Results indicate increased expression of TREM2 in ALS patient samples as compared to non-neurological controls. **c,** Endogenous TDP-43 of frozen autopsied spinal cord specimens from ALS patients was co-immunoprecipitated with TREM2. Experiments were independently repeated three times. **d-g,** Cartoon models of the human TREM2 extracellular ligand-binding domain in complex with four low-complexity domain (LCD) fragments of TDP-43. **h,** Surface model of the TREM2 complex showing overlapping binding sites of GFN and NFG and distinct binding sites of NFG, AMM, and SWG. (NFG: N^312^FGAFS^317^; AMM: A^321^MMAAA^326^; SWG: S^333^WGMMGMLASQ^343^; GFN: G^396^FNGGFG^402^). **i**, SPR analysis of the binding between recombinant hTREM2 ECD (Met1-Ser174) with full length hTDP43. hTREM2 ECD was immobilized on a CM5 BIAcore chip and interacted with hTDP43 at the indicated concentrations. **j**, Recombinant hTREM2 ECD (Met1-Ser174) was immobilized on a CM5 BIAcore chip and binding of SWG and GFN at the indicated concentrations were determined by SPR. Data represented as mean ± SEM. Significances were calculated using student t test; *n.s.*, not significant; \**P* < 0.05, \*\**P* < 0.01, *** *P* < 0.001.

### TREM2 interacts with residues 333–402 of hTDP-43 *in silico* and *in vitro*

To further investigate the interaction between TREM2 and hTDP-43 at the atomic level, we performed a computational analysis. Our *in vitro* study already showed that TREM2 interacted with residues 216-414 of hTDP-43. Therefore, we carried out microsecond molecular dynamics (MD) simulations using forcefield FF12MC and the crystal structures of the extracellular ligand-binding domain (ECD) of hTREM2 and low-complexity domain (LCD) fragments of hTDP-43 ^33, 34^. The four LCD fragments were N^312^FGAFS^317^ (NFG), A^321^MMAAA^326^ (AMM), S^333^WGMMGMLASQ^343^ (SWG), and G^396^FNGGFG^402^ (GFN) (**Fig. 7d-g**). They were chosen because of their involvement in irreversible stress granules ^34^. Our visual inspection suggested the structural fitness between the residues in complementarity-determining regions (CDRs) of the TREM2 ECD and those in four LCD fragments of hTDP-43.

Four sets of 20 distinct, independent, unrestricted, unbiased and isobaric–isothermal MD simulations were carried out for each of the four LCD fragments in complex with the hTREM2 ECD. These fragments were manually docked into the same ECD region to have the LCD fragment surrounded by CDR2 on one side and CDR1 and CDR3 on the other side. Interestingly, the simulations showed that AMM and SWG moved out of the CDR1–3 to interact with β-strands residues 75–78 and 65–70 of the ECD, respectively (**Fig. 7e, f**). The relative interaction energies of the hTREM2 ECD with SWG, GFN, NFG, and AMM were -74, -38, -38, and -33 kcal/mol, respectively. More interestingly, GFN and NFG remained in the CDR1–3, and NFG (but not GFN) moved closer to R47 of the ECD with its two phenyl rings interacting with W44 and W70 of the ECD (**Fig. 7d, g**). There was an overlap for the binding sites on the ECD for NFG and GFN, but the binding sites for NFG, AMM, and SWG were distinct (**Fig. 7h**). These results suggest that TREM2 is in contact with the LCD residues 312–343 that encompass NFG, AMM, and SWG and constitute a potential epitope of hTDP-43 for TREM2, whereas GFN represent another epitope comprising the LCD residues 333–402 that encompass SWG and GFN. In addition, our simulations revealed that none of the four LCD fragments interact with the two glycosylation sites (N20 and N79) of the ECD domain of TREM2. Therefore, the interaction between TREM2 and hTDP-43 is likely independent of the glycosylation state of the ECD domain.

We then performed the Biacore-based surface plasmon resonance (SPR) analyses to study the direct interactions of hTREM2 ECD with four LCD fragments or the full-length hTDP-43. According to these analyses, hTREM2 indeed interacted with GFN, SWG, and the full-length hTDP-43 in a dose-dependent manner with *K*_D_ values of 1.86 × 10^-4^ M, 1.33 × 10^-5^ M, and 1.18 × 10^-7^ M, respectively (**Fig. 7i, j**). However, we did not observe the interaction of hTREM2 with NFG or AMM under the same conditions (**Extended Data Fig. 10b**), possibly due to the interference of the hTREM2 immobilization involving K42, K57, and/or K123 with the binding of NFG or AMM. Nevertheless, these SPR results are consistent with the interactions of hTREM2 with SWG and GFN predicted by the MD simulations. Therefore, the SPR results support the direct interaction of hTREM2 with hTDP-43 that was inferred from our *in vivo*, *in vitro* and *in silico* studies.

## DISCUSSION

A major challenge to ALS research is identifying the intrinsic mechanisms that can sense an aberrant CNS environment during the initial stage of disease, and protect motor neurons from degeneration ^35^. Our study provides a comprehensive analysis of the impact of TREM2 deficiency on microglial phenotypic and functional change in mouse models of TDP-43 proteinopathy. These results demonstrate that TREM2 regulates the microglial response to, and phagocytic clearance of, pathological TDP-43, thus reducing TDP-43 proteinopathy and neurodegeneration. We further reveal that TDP-43 interacts with TREM2 *in silico*, *in vitro* and *in vivo*. This explains why mice with TREM2 deficiency exhibit reduced microglial clearance of TDP-43 and worse motor deficits. Our study suggests a protective role of microglial TREM2 in TDP-43 clearance to alleviate TDP-43 neurodegeneration.

Here we used the neonatal hTDP-43 overexpression model which displayed a progressive TDP-43 pathology associated with motor neuron degeneration, replicating primary motor symptoms of ALS. However, this model has its own limitations and does not fully recapitulate clinical pathology. First, neonatal model started to express hTDP-43 at early age while the risk of developing ALS peaks at 50 to 75 years of age for patients ^36^; Secondly, neonatal model widely expressed hTDP-43 in all CNS regions at the same time, while TDP-43 pathology spreads in ALS patients following a certain progression pattern, with pTDP-43 accumulation in upper motor neurons in the cortex, lower brainstem and lower motor neurons in the spinal cord at the initial stage ^37^; Thirdly, neonatal model developed accumulated TDP-43 pathology and exhibited motor deficit in a short period time after AAV injection, while the disease progression and survival are highly variable in clinics ^36^. Fourthly, neonatal model is a hTDP-43 overexpression model, while the typical TDP-43 pathology observed in clinic are the mislocalization of TDP-43 from the nucleus to the cytoplasm and formation of phosphorylated and ubiquitinated cytosolic inclusions ^8,38, 39^. To partially overcome the limitations, we also used adult local injection model which developed TDP-43 pathology only in the primary motor cortex as well as a transgenic rNLS8 mouse model in which hTDP-43 accumulated in cytoplasm after expression.

### TREM2 mediates microglial activation in the course of TDP-43 pathology

The intrinsic heterogeneity of microglia allows them to acquire multiple phenotypes in response to extrinsic stimuli ^40, 41^. Our study provides strong evidence of phenotypic impact of TREM2 on microglia response to TDP-43 pathology. Two recent studies have demonstrated that TREM2 deficiency attenuated microglial responses to Aβ and tau pathologies ^42, 43^. In contrast, elevated TREM2 gene dosage conferred a rescuing effect in AD mice by altering microglial morphology and reprogramming microglia responsivity ^44^. Here we found that TREM2 deficiency significantly blocks the microglial activation in response to TDP-43 accumulation. At least two potential mechanisms could underlie TREM2-dependent microglia activation: (1) The direct interaction with pathological TDP-43 activates TREM2 signaling pathway via DNAX-activation protein 12 (DAP12) and phosphorylation of Syk ^45^. (2) Neuronal contact-dependent activation is another possibility. This is supported by increased number of microglia-neuron physical interaction in response to TDP-43 pathology. In addition, prior study showed that lack of TREM2 impairs microglial migration towards damaged neurons ^46^. Thus, one can presume that TDP-43 damaged neurons express signals sensed by TREM2 that might trigger microglial activation. Together, our study shed new light on the “find me” mechanisms that underline TREM2-mediated microglial activation in response to TDP-43 related neurodegeneration.

The next important question to address is what molecular changes occur during microglia phenotypic reprogramming after sensing stimuli. We showed TREM2 deficiency prevents the downregulation of two microglia homeostatic markers P2Y12 and TMEM119 in the context of TDP-43 related neurodegeneration. In addition, microglial morphology was locked into homeostatic states in TREM2 KO mice when compared with WT mice in response to TDP-43. However, how these molecular changes can remodel the cytoskeleton and morphology, underlying phenotypic change, remains unknown. In the context of AD mice, brain transcriptomic analysis demonstrated that elevated TREM2 gene dosage upregulated reactive microglia genes and dampened multiple disease-associated microglia genes ^44^. Therefore, it is likely that the activation of microglial TREM2 by TDP-43 may regulate the transcription factors that affect homeostatic gene expression in microglia. Future comprehensive transcriptome-wide and network analysis studies are needed to investigate the potential molecular candidates that could be tapped to regulate microglia activation in the course of TDP-43 pathology.

Interestingly, we found that TREM2 deficiency did not diminish the expansion of microglial numbers in our disease model, which is reminiscent of other microglia receptors involved in proliferation. Previous study demonstrated that TREM2 is important for microglial proliferation in response to KA treatment and promotes microglia survival via Wnt/β-Catenin pathway ^47^. Therefore, it is likely that TREM2 may promote microglial proliferation in a context-dependent manner. These results also raised an important question as to the major driver for microglia proliferation in response to TDP-43. Previous studies have reported that colony–stimulating factor-1 receptor (CSF1R) is critical in microglial proliferation in response to peripheral nerve injury and in seizures ^48, 49^. In addition, TREM2 and CSF1R share the similar adaptor protein DAP12 and downstream signals ^50^. Thus, it would be interesting to explore whether CSF1R mediated pathway can independently serve as a main regulator for microglial proliferation in TDP-43-related neurodegeneration.

### TREM2 mediates microglial phagocytic clearance of pathological hTDP-43

Our study demonstrates that TREM2 can directly mediate microglia phagocytosis of pathological TDP-43 proteins. The presence of hTDP-43 in microglia is unlikely due to the expression of AAV-hTDP-43. AAV virus is known to have very limited ability to infect microglia ^51^. Instead, we provided strong evidence that microglia actively phagocytosed pathological hTDP-43 in WT mice which was significantly attenuated in TREM2 KO mice. This is consistent with observations in AD mouse models in which TREM2 mediated microglial phagocytosis can limit amyloid plaque growth ^42^. Phagocytosis describes the cellular process of eating, including receptor mediated recognition, engulfment, and degradation of particulate organisms or structures with size over 0.5 µm ^52^. As professional phagocytes, microglia orchestrate innate brain immune response by eliminating detrimental structures ^52, 53^. Typical microglia phagocytosis receptors include Toll-like receptors (TLRs) and TREM2. The former has high affinity to bind pathogen-associated molecular patterns (PAMPS), and the latter can recognize apoptotic cellular substances ^54^. In addition, other receptors like complement receptors (CR3), pyrimidinergic receptors (P2Y6) and TAM receptors are also involved in microglia phagocytosis ^55–57^. Although our results strongly suggest that microglial TREM2 is a key receptor in phagocytosis of TDP-43, it would be of interest to explore whether other microglia phagocytosis receptors could work together with TREM2.

Our study sheds novel light on the antigen presenting function of microglia related to the phagocytosis of TDP-43. Microglia have been reported to serve as antigen-presenting cells in response to inflammatory challenge ^58^ and to other neurodegenerative diseases, like AD and PD ^59, 60^. After uptake of the pathological TDP-43 proteins, the further degradation of TDP-43 likely leads to the antigen presentation. We found that microglia do not express CD11c under normal conditions, which is in agreement with the poor antigen presenting ability of microglia in general ^61^. However, we identified a distinct microglia subpopulation that highly expressed CD11c in response to hTDP-43 expression in WT mice. This CD11c^+^ subpopulation of microglia is dependent on TREM2 and is highly phagocytic towards hTDP-43. In addition, considering the function of CD11c in antigen-presenting cells, these observations raised the question of whether CD11c^+^ microglia could serve as those cells during TDP-43 related neurodegeneration. Indeed, increased CD4^+^ T cell accumulation has been reported in patient with ALS ^62^ and other chronic neurodegeneration pathologies ^63^. Moreover, T cell related immunotherapy has been applied in clinical trial and brings new hope for ALS patients ^64^. These lines of evidence warrant future studies on the interaction of the CD11c^+^ microglia and infiltrated T cell in TDP-43 related neurodegeneration.

Importantly, our data showed that TDP-43 induced CD11c^+^ upregulation is dependent on TREM2, as CD11c^+^ microglia are absent in TREM2 KO mice. We further demonstrated the beneficial function of CD11c^+^ microglia in clearance of pathological TDP-43 proteins. These findings are consistent with an AD mouse model where strong upregulation of microglial CD11c was significantly attenuated by TREM2 deficiency ^5, 65^. Single-cell RNA analysis of CNS immune cells in AD, ALS and aging revealed a TREM2 mediated subset disease-associated microglia (DAM) with unique transcriptional signature^65^. This subpopulation was further demonstrated to play a neuroprotective role and is characterized by a downregulation of homeostatic gene signature and upregulation of genes involved in microglia phagocytosis and lipid metabolism. Interestingly, CD11c is also high expression in DAM. The CD11c^+^ microglia are localized near amyloid plaques, potentially to counteract amyloid deposition ^23^. In other disease models, like a cuprizone-induced demyelination model, CD11c^+^ microglia can accelerate the repair of damaged white matter ^66^. Our results of TREM2-dependent CD11c expression in response to TDP-43 raised these intriguing questions: (1) how TREM2 regulates microglial CD11c expression, (2) the function of CD11c^+^ microglia subpopulation during TDP-43 pathology, and (3) the unique signatures of the CD11c^+^ microglia. Previous transcriptional analysis of CD11c^+^ microglia accumulating around amyloid plaques revealed a higher expression of TREM2 in a mouse model of Alzheimer’s disease ^23^. Whether TREM2 is upregulated in CD11c^+^ microglia subpopulation in the context of TDP43 pathology needs further investigation. We are cognizant of the fact that hTDP-43 overexpression model cannot perfectly mimic ALS patient pathology. Nevertheless, our study still provides insight into the molecular targets of reprogramming microglia via TREM2 in sensing and regulating TDP-43-related neurodegeneration.

### TDP-43 interacts with TREM2

It has been reported that potential ligands for TREM2 are lipids, APOE, and Aβ ^5, 6, 67^. Our present results demonstrate that TDP-43 also interacts with TREM2 both *in vitro* and *in vivo* and thus may act as a ligand for TREM2. In addition, our computational simulation provides the potential TDP-43 binding sites on TREM2. This interaction between TREM2 and TDP-43 may represent an intriguing mechanism for microglial TREM2 mediated sensing and clearance of pathological TDP-43 protein. Previous studies have shown that TDP-43 can be released by extracellular vesicles and then taken up by other neurons and contribute to neuronal damage ^68^. It is also possible that TDP-43 can be released into the extracellular space as free cargos by unconventional protein secretion pathways which are induced by cellular stress. Indeed, we are able to detect extracellular hTDP-43 in *in vitro* cultures of iPSC neurons. Therefore, neuronal TDP-43 accumulation may lead to its release, which can be sensed and phagocytosed via microglial TREM2.

Colocalization of TREM2 with TDP-43 by immunostaining provided the evidence of TREM2 interaction with TDP-43. Interestingly, our TREM2 staining revealed an intra microglial TREM2/TDP-43 complex, indicating a possible function of sTREM2 in mediating microglia phagocytic clearance of TDP-43. Canonical TREM2 undergoes sequential proteolytic processing by a disintegrin and metalloprotease (ADAM) and γ-secretase to generate sTREM2 ^69, 70^. After cleavage, sTREM2 can be released into the extracellular space and physically interacts with amyloid plaques ^71^. A recent functional study has further revealed that sTREM2 is able to enhance microglial phagocytosis and degradation of Aβ by promoting microglia proliferation and migration to the vicinity of the plaques ^72^. Hence, in the context of TDP-43 pathology, sTREM2 may recognize the extracellular TDP-43 and recruit microglia for phagocytic clearance. However, further studies are needed to test this hypothesis.

TREM2-mediated clearance mechanism is supported by structural analysis showing that neurodegenerative disease related mutations in TREM2 affects its ligand binding ability ^73^. Indeed, AD-associated mutations compromise interactions of TREM2 with Aβ ^6^ and GLU ^74^. Based on our *in vitro* study which showed hTREM2 interacted with residue 216-414 of hTDP-43, and given the limitations of current computing technologies and the intrinsic instability of hTDP-43, we performed fragment-based MD simulations instead of MD simulations of TREM2 in complex with a full-length TDP-43 model with statistical relevance. Without any bias from disease-associated mutations in TREM2 and hTDP-43, our fragment-based MD simulations suggest that TREM2 is in contact with two potential epitopes of hTDP-43 (residues 312–343 or 333–402). Subsequent SPR analysis has confirmed the 333–402 epitope, although the confirmatory SPR data for the 312–343 epitope are challenging to obtain due to the hTREM2 immobilization involving surface lysines. Interestingly, the NFG fragment of the 312–343 epitope and the SWG fragment of both epitopes contain many mutations of the hTDP-43 in ALS patients. Consistently, there are correlations among mutations in hTDP-43 and/or hTREM2 and the propensity of hTDP-43 for aggregation ^75^. While further studies such as in situ cross-linking are needed to biochemically confirm the hTREM2 interactions with residues 312–343 or 333–402 of hTDP-43 *in vivo*, our present immunoprecipitation, SPR, and MD simulation data offer new insight into the potential interaction site of hTREM2 and hTDP-43, highlighting that hTDP-43 is an intriguing new ligand for microglial hTREM2. In summary, our study found that TREM2-mediated microglial activation is an early protective response to TDP-43. The underlying mechanism may involve the interaction between TDP-43 and TREM2 that mediated phagocytosis of TDP-43 by CD11c^+^ microglia subpopulation. These findings point out microglial TREM2 as a potential therapeutic target for ameliorating TDP-43-related neurodegeneration.

## Methods

### Mice

The TREM2 knockout (KO) mouse strain was kindly provided by Dr. Marco Colonna at the Washington University School of Medicine, St. Louis. Wild-type (WT) C57BL/6J mouse strain, CX3CR1^GFP/+^ mice and rNLS8 ^22^ were purchased from Jackson Labs. Mouse lines were housed with littermates with free access to food and water on a 12 hour light/dark cycle. All animal procedures, including husbandry, were performed under the guidelines set forth by the Mayo Clinic Institutional Animal Care and Use Committee.

### Human samples

Post-mortem brain samples were dissected from the frozen brains of 16 ALS cases (age 67 ± 9.4 years, mean ± s.d.) and 12 controls (age 74.7 ± 10.6 years) from the Brain Bank for Neurodegenerative Disorders at Mayo Clinic Jacksonville. The study was approved by Mayo Biospecimen Committee. For clinic pathological studies, cases were included only if they had good quality medical documentation and there was diagnostic concurrence of at least minimum of two neurologists. A summary of cases, including clinical and pathologic analyses, are listed in supplementary Table 2.

### Cell culture and transfection

Human embryonic kidney (HEK) 293T cells were cultured in DMEM (Gibco, Thermo Fisher Scientific) supplemented with 10% FBS (Gibco, Thermo Scientific). Cells were cultured at 37°C with 5% CO_2_. HEK293 cells were co-transfected with plasmids encoding wide-type myc-human TREM2 and GFP-human TDP-43 (residues 216-414, Addgene, 28197) or GFP control by the calcium phosphate precipitation method. The expression of TREM2 or TDP-43 was assessed 24-72 h after infection.

Induced iPSC cultures were prepared as previously reported ^76^. 6-well plates were coated with hESC Qualified Geltrex (Gibco, ThermoFisher) and incubated at 37°C and 5% CO_2_ for at least 1h. Differentiated iPSC neurons were cultured in media composed of KO DMEM/F-12, 2% B27 supplement (Gibco, ThermoFisher), 100 U/mL penicillin, 100 lg/mL streptomycin, 1% Glutamax, 1 mmol/L dibutyryl cyclic-adenosine monophosphate (cAMP; Sigma), 5 lg/mL plasmocin (Invivogen), 1 mmol/L sodium pyruvate and 10 ng/mL each of BDNF (PreproTech), IGF1 (Corning) and GDNF (PeproTech). To express proteins in iPSC neurons, 1 µL each of AAV9.CAG.hTDP-43-GFP (9.0E+13 viral genomes/mL; AAV9 empty vector as control) and AAV1.Tdtomato (5.0E+12 viral genomes /mL; Addgene, 59462) were added to 1 mL culture medium.

### Intracerebroventricular injections

AAV was injected intracerebroventricularly (ICV) into C57BL/6 mouse or TREM2 KO mouse pups on postnatal day 0 (P0). ICV injections were performed as described ^77^. Newborn mice were put onto a cold metal plate for 2-3 mins to induce hypothermic anesthesia. Cryoanesthetized neonates were injected using 10 uL Hamilton syringes affixed with a 32-gauge needle at an angle of 45 degree to a depth of 3 mm. The injection site was located at 2/5 of the distance from the lambda suture to each eye and 2 uL of virus (9.0E+12 viral genomes/mL) was slowly injected into each ventricle. After injection, pups were allowed to completely recover on a warming blanket and then returned to the home cage.

### Stereotaxic intracerebral injection

Adult wildtype C57BL/6 mice or TREM2 KO mice at 2 months of age were anesthetized by 2% isoflurane. 0.2 μL AAV (9.0E+12 viral genomes/mL) was then injected into the right primary motor cortex at the following stereotaxic coordinates: 1.4 mm anterior to bregma, 1.5 mm lateral to the midline, 0.5 mm ventral to the dura, with bregma set at zero. The microinjections were carried out at a rate of 0.04 μL/min, using an automated stereotaxic injection apparatus (Model UMC4, World Precision Instruments Inc.). The microsyringe was left in place for an additional 10 mins after each injection.

### Immunofluorescence staining

Tissue was prepared as previously reported ^78^. Mice were anesthetized with isoflurane (5% in O_2_) and then intracardially perfused with 40 ml cold PBS, followed by 40 ml cold 4% paraformaldehyde (PFA). Brains were removed and post-fixed in 4% PFA for an additional 6 hours, then transferred to 30% sucrose in PBS for three days. 20 μm slices were cut with a cryostat (Leica). For immunofluorescence staining, the sections were blocked for 60 mins with 10% goat serum in TBS buffer containing 0.4% Triton X-100 (Sigma), and then incubated overnight at 4 °C with a primary IgG antibody: rabbit anti-NeuN (1:500, Abcam, ab104225), rabbit anti-CD68 (1:500, Abcam, ab125212 ), hamster anti-CD11c (1:200, Thermo, 14011482), rabbit anti-P2Y12 (1:400, Anaspec, AS-55043A), rabbit anti-Iba1 (1:500, Abcam, ab178847), rabbit anti-p-TDP-43 (1:200, Proteintech, 22309-1-AP), mouse anti-GFAP (1:500, CST, 3670), rabbit anti-CNPase (1:500, Proteintech, 13427-1-AP), rabbit anti-TMEM119 (1:500, Abcam, ab209064) or sheep anti-TREM2 (1:50, R&D AF1729). After three washes with TBST, sections were exposed to appropriate secondary antibody for 60 mins at room temperature (1:500, Alexa Fluor goat anti-rabbit 594, Thermo, a32740,1:500, Alexa Fluor goat anti-ArHm 647, Abcam, ab173004 or 1:500, Alexa Fluor donkey anti-sheep 594, Thermo, a11016), washed and mounted and coverslipped with Fluoromount-G (SouthernBiotech). Fluorescent images were captured with a confocal microscope (LSM510, Zeiss), and cells of interest were counted and fluorescence signal intensity was quantified using ImageJ software (National Institutes of Health, Bethesda, MD).

### Immunohistochemistry

Free floating brain sections (30 μm) were rinsed 3 times in TBS, and incubated in 3% H_2_O_2_ for 30 minutes to quench endogenous peroxidase activity. IHC staining for hTDP43 antibody (mouse, 1:500, Proteintech, 60019-2-Ig) and p-hTDP43 antibody (mouse, 1:500, Proteintech, 22309-1-AP) were performed using M.O.M. immunodetection kit (Vector Laboratories BMK-2202) following the manufacturer’s instructions. After primary antibody (1:500), biotinylated appropriate secondary antibodies were added for 2 hours at room temperature. Vectastain-ABC Kit (Vector Laboratories) and ImmPACT DAB substrate peroxidase (HRP) kit (Vector Laboratories) were used to amplify and detect signals. Subsequently, sections were washed, mounted, and dehydrated through 70%, 95%, and 100% ethanol solutions. After immersing in xylene, coverslips were placed on slides using DEPEX mounting medium (Electron Microscopy Science) for observation.

### Nissl staining

Sample slices were incubated in 100% ethanol for 6 mins, and then defatted in Xylene for 15 mins, followed by another 10 mins in 100% ethanol. After rinsing in distilled water, the slides were stained with 0.5% cresyl violet acetate for 15 mins, and again rinsed in distilled water. The sections were then placed in differentiation buffer (0.2% acetic acid in 95% ethanol) for 2 mins, dehydrated with ethanol and Xylene, and then mounted with Depex medium. Neuronal number in the layer 5 of primary motor cortex was quantified following previous established standard criteria: (1) pyramidal neurons exhibited a characteristic triangular shape with a single large apical dendrite extending vertically towards the pial surface, (2) non-pyramidal neurons were identified by the absence of the preceding criteria, by the presence of two or more small calibre processes, and by their generally smaller size ^79^.

### Mass cytometry/CyTOF of isolated brain cells

Microglia antibody panel based on surface markers (supplementary Table1) was designed for brain mass cytometry/cytometry by time of flight (CyTOF). All the antibodies were provided by Mayo CyTOF core (Rochester, MN). The whole mouse brain was dissociated into a single-cell suspension using brain tissue dissociation kits following the manufacturer’s instructions (Adult Brain Dissociation Kit, Miltenyi Biotec, 130-107-677). Collected cells were then incubated with metal-conjugated antibodies in cell staining buffer (ProductMaxpar Cell Staining Buffer, Fluidigm, CA). The barcoded sample was loaded onto a Helios CyTOF® system (Fluidigm) using an attached autosampler and were acquired at a rate of 200-400 events per second. Data were collected as .FCS files using the CyTOF software (Version 6.7.1014). After acquisition intrafile signal drift was normalized to the acquired calibration bead signal using the CyTOF software. Cleanup of cell debris, removal of doublets and dead cells was performed using FlowJo software version 10.5.3 (Ashland, OR). Cleaned FCS files were analyzed by Cytobank (Santa Clara, CA) and FlowJo.

### Isolation of RIPA-soluble and sarkosyl-insoluble proteins

Mouse brain tissue was lysed in RIPA lysis buffer (Cell signaling), supplemented with protease inhibitors cocktail and phosphor inhibitor (Thermo Fisher Scientific), and kept on ice for 30 mins. The brain homogenate was then ultracentrifuged at 50,000g for 15 mins at 4°C. The supernatant was transferred to a new tube as RIPA-soluble protein fraction. The pellet was homogenized in 10% sucrose buffer before being ultracentrifuged at 50,000g for 15 mins at 4°C. After ultracentrifuge, the pellet was discarded and the supernatant was transferred to a new tube and incubated with sarkosyl (Sodium lauroyl sarcosinate) at a final concentration of 1% for 1 hour at 37°C. Following incubation, the samples were ultracentrifuged at 50,000g for 30 mins at 4°C. The supernatant was discarded and the pellet was re-suspended to obtain the insoluble protein fraction. Soluble and insoluble fractions were then boiled in SDS-PAGE loading buffer for immunoblotting. Blots were probed with antibodies for mouse anti-hTDP-43 (1:1000, Proteintech, 60019-2-Ig), rabbit anti-total TDP-43 (1:1000, Proteintech, 10782-2-AP), mouse anti-human TDP-43, phospho Ser409/410 (1:2000, COSMOBIO, CAC-TIP-PTD-M01), rabbit anti-GFP (1:1000, Cell signaling, 2956s) and mouse anti-GAPDH (1:5000, Santa Cruz, sc-32233).

### Immunoprecipitation assay

For *in vitro* binding assays, we transfected HEK 293T cell with myc tagged TREM2 and GFP-hTDP-43 or GFP control plasmids. Cell lysates were prepared in IP lysis buffer (50 mM Tris-HCl, pH 7.5, 0.5% Nonidet P-40, 150 Mm NaCl, 5Mm EDTA) supplemented with protease inhibitors cocktail and phosphotase inhibitor (Thermo Fisher Scientific), and kept on ice for 30 mins. The lysates were then centrifuged at 14,000g for 15 mins at 4°C. The supernatant protein concentration was measured and normalized between samples. Myc-Trap agarose beads were used to pull down the Myc-TREM2 protein following the manufacturer’s instructions (Chromotek, yta-10). GFP-Trap agarose beads were used to pull down the GFP-human TDP-43 (residues 216-414) or GFP protein (Chromotek, gta-10). Blots were probed with antibodies for mouse anti-Myc antibody (1:1000, Sigma, SAB4700447), rabbit anti-TDP-43 (1:1000, Proteintech, 10782-2-AP), rabbit anti-GFP (1:1000, Cell signaling, 2956s).

For *in vivo* binding assays, TDP-43/TREM2 interaction was detected in rNLS8 mouse with 4 weeks DOX off and 2 weeks DOX on. Mouse TREM2 was immunoprecipitated from 400-500 ug protein per sample using 1.5 ug mouse TREM2 biotinylated antibody (R&D, BAF1729). After overnight incubation at 4°C, biotinylated antibody was pulled down using NeutrAvidin Agarose beads (Thermo Fisher 29200) for at least 2 h at 4°C. Avidin beads were collected by centrifugation at 2,000 ×g for 3 mins and washed four times with IP lysis buffer, and then boiled in SDS-PAGE loading buffer for immunoblotting. Blots were probed with antibodies for rat anti-mouse TREM2 antibody [5F4] (1:500, abcam, ab252876) and mouse anti-hTDP-43 (1:1000, Proteintech, 60019-2-Ig).

Endogenous TDP-43/TREM2 interactions were also determined using frozen autopsied specimens of ALS patients’ spinal cord. Lysates containing 400-500 μg protein were incubated with rabbit anti-human TREM2 antibody (Cell Signaling, 91068) / Dynabeads protein A (Thermo fisher, 22810) at 4°C overnight. Protein A beads were collected by centrifugation at 2,000 ×g for 3 mins and washed four times with IP lysis buffer, and then boiled in SDS-PAGE loading buffer for immunoblotting. Blots were probed with antibodies for goat anti-human TREM2 antibody, (1:1000, R&D, AF1828) and mouse anti-h-TDP-43 (1:1000, Proteintech, 60019-2-Ig).

### Immunoblotting

Mouse brain tissue or human tissue samples were lysed in RIPA lysis buffer (Cell signaling), supplemented with protease inhibitors cocktail and phosphatase inhibitor (Thermo Fisher Scientific), and kept on ice for 30 mins. The lysates were then centrifuged at 14,000 ×g for 30 mins at 4°C. The supernatant protein concentration was measured and normalized between samples, and was boiled in SDS loading buffer. Prepared samples were subjected to 4%–20% SDS-PAGE (Bio-Rad laboratories).

### Liquid Chromatography with tandem mass spectrometry (LC-MS/MS) and affinity purification coupled with mass spectrometry (AP-MS) interactome analysis

To detect interacting proteins of TREM2, HEK 293 cells or rNLS mice brains were lysed in lysis buffer (50 mM Tris-HCl, protease inhibitors cocktail (Thermo Fisher Scientific), phosphatase inhibitor (Thermo Fisher Scientific), and 0.002% zwittergent 3-16) and kept on ice for additional 30 mins. After centrifugation, the supernatant were incubated with rabbit anti-human TREM2 antibody (Cell Signaling, 91068) or anti-mouse TREM2 antibody on a rotator, respectively. After 3 washes, immunoprecipitated proteins were eluted by boiling in Lammaeli buffer for immunoblotting. Alternatively, immunoprecipitated proteins were eluted by boiling in 1% SDS, resolved by non-reducing gel and subjected to reduction, alkylation and trypsin digestion using an in-gel digestion protocol. The extracted tryptic peptides were dried in a speed vac. The dried peptides were reconstituted in sample loading buffer (0.2% formic acid, 0.1% TFA, 0.002% zwittergent 3-16) and analyzed by nano-ESI-LC/MS/MS with a Q Exactive mass spectrometer coupled to a dionex nano-LC system (Thermo Scientific; Waltham, MA) with a 32 cm, 100/15 um, Acclaim 2.2 um column. The LC system used a gradient with solvent A (2% ACN, 0.2% formic acid, in water) and solvent B (80% ACN, 10% IPA, 0.2% formic acid, in water) with a 400 nL/min flow rate as follows: 4-5 min 5% B; 5-95 min 5-45 % linear gradient; 95-98 min 45-95% linear gradient; 98-102 min 95% B; 102-104 min 95-10% B linear gradient; 104-107 min 10% B; 107-115 min 10-95% B; 115-118min 95% B; 118-121 min 95-5% B linear gradient; 121-127 min 5% B. Mass spectrometer was operated in data-dependent mode with a MS survey scan (350-1800 m/z) in Orbitrap (70,000 resolution 200 m/z, 3 × 106 AGC target and 100 ms maximal ion time) and 20 MS/MS fragments scans (17,500 resolution, 2 × 105 AGC target, 50 ms maximal ion time, 28 normalized collision energy, 2.5 m/z isolation window) with a dynamic range set to 25 seconds. The tandem mass spectra were searched using SEQUEST ^80^ search algorithms against a Uniprot database (human 2019_07 release for 293T CoIP and mouse 2019_08 release supplemented with human TDP-43 protein sequence for mouse CoIP) through the Proteome Discoverer platform (version 2.5, Thermo Scientific). The search parameters included a maximum of two missed cleavages; carbamidomethylation at cysteine as a fixed modification; N-terminal acetylation, deamidation at asparagine and glutamine, oxidation at methionine as variable modifications; the monoisotopic peptide tolerance, 5 ppm and MS/MS tolerance, 0.02 Da. The target-decoy analysis was used for validation. The false discovery rate (FDR) was set to 1% at both peptide and protein level.

### Stable isotope labeling with amino acids in cell culture (SILAC)

Two sets of HEK293T cells were cultured in DMEM containing “light” (^12^C_6_-arginine and ^12^C_6_-lysine) or “heavy” (^13^C_6_-arginine and ^13^C_6_-lysine) amino acid supplemented with FBS, penicillin and streptomycin, separately. The cells gown in light medium were transfected with empty vector while the cells grown in heavy medium were transfected with hTREM2 plasmid. Forty-eight hours after transfection, both sets of cells were lysed in lysis buffer (50 mM Tris pH 7.4, 150 mM NaCl, 1% n-Octylglucoside) in the presence of cocktail protease and phosphatase inhibitors. The protein concentration of the lysates from light and heavy cells were measured by BCA method. Equal amounts of light and heavy lysates were subjected to immunoprecipitation by anti-hTREM2 antibody. After washing, the immunoprecipitates from light and heavy lysates were mixed and subjected to a trypsin digestion protocol on S-Trap according to the manufacturer’s instruction (ProtiFi). The resulting peptides were analyzed by LC-MS/MS on a high-resolution orbitrap-equipped mass spectrometer. The acquired mass spectra were analyzed on the MaxQuant proteomics analysis platform. The lack of light peptide in the mass spectrum will indicate a true TREM2 interacting protein.

### Hindlimb clasping

To examine hindlimb clasping, mice were gently removed from their home cage and suspended by the tail for 10 s and provided a hindlimb clasping score. If the mouse exhibits normal escape extension without limb clasping, it was assigned a score of 4. If one hindlimb exhibits incomplete splay and loss of mobility and toes exhibit normal splay, it received a score of 3. If both hindlimbs exhibit incomplete splay and loss of mobility and toes exhibit normal splay, it received a score of 2. If both hindlimbs exhibit clasping with curled toes and immobility, it received a score of 1. If forelimbs and hindlimbs exhibit clasping and are crossed, curled toes and immobility, it received a score of 0.

### Open field testing

Locomotor activity of mice was assessed in sound-insulated, rectangular activity chambers with continually running fans, infrared lasers, and sensors (Med Associates, St Albans, VT, USA). Prior to testing, mice were acclimatized to the room for 1 h. Mice were habituated for 5 mins in the Open Field chamber (without recording) then placed back to home cage for another 5 mins. Afterwards, mice were introduced back to the chambers and activities were monitored for 30 mins. Locomotor function was quantified by the total distance mice travelled in the chamber which was recorded on a computer with Med-PC software Version 4.0.

### Rotarod

The Rotarod performance test evaluates mouse balance and motor coordination. Mice were brought to the test room 1 h before testing. Rotarod tests were performed using a five-lane Rotarod apparatus (Med Associates Inc), started at 4 revolutions per minute and steadily accelerated to 40 revolutions per minute over a 5 mins period. Each mouse was tested 3 times at 10-min intervals.

### Molecular dynamics (MD) simulations

The LCD fragment was manually docked into the paratope region of the crystal structure of the ECD (Residues 21-219) of human TREM2 (PDB ID: 6B8O) as described above. The resulting complex was neutralized with counterions and then energy-minimized according to a protocol described below. In this complex, all ionic residues were treated as AMBER residues of ASP, GLU, ARG and LYS, all histidine residues were treated as HIE, and all cysteine residues were treated as CYX. The topology and coordinate files of the complex were generated by the tLeAP module of AmberTools 13 (University of California, San Francisco). The energy minimization and MD simulations were performed using SANDER and PMEMD of AMBER 16 (University of California, San Francisco) with force field FF12MC ^33^. The energy minimization used dielectric constant of 1.0 and 100 cycles of steepest-descent minimization followed by 900 cycles of conjugate-gradient minimization. The energy-minimized model was solvated with the TIP3P water using “solvate box m TIP3PBOX 8.2” and energy minimized for 100 cycles of steepest-descent minimization followed by 900 cycles of conjugate-gradient minimization using the SANDER module ^81^. The resulting system was heated from 5 K to 320 K at a rate of 10 K/ps under constant temperature and constant volume, equilibrated for 10^6^ time steps under constant temperature of 320 K and constant pressure of 1 atm employing isotropic molecule-based scaling, and finally simulated, without any harmonic restraint, in 20 316-ns, distinct, independent, unrestricted, unbiased, and isobaric–isothermal MD simulations with FF12MC using the PMEMD module of AMBER 16 with a periodic boundary condition at 320 K and 1 atm. The 20 unique seed numbers for initial velocities of Simulations 1–20 were taken from Ref. ^82^. All simulations used (i) a dielectric constant of 1.0, (ii) Langevin thermostat ^83^ with a collision frequency of 2 ps^-1^, (iii) the Particle Mesh Ewald method to calculate electrostatic interactions of two atoms at a separation of >8 Å ^84^, (iv) Δt = 1.00 fs of the standard-mass time ^33, 85^, (v) the SHAKE-bond-length constraints applied to all bonds involving hydrogen, (vi) a protocol to save the image closest to the middle of the “primary box” to the restart and trajectory files, (vii) the revised alkali and halide ions parameters ^86^, (viii) a cutoff of 8.0 Å for nonbonded interactions, (ix) the atomic masses of the entire simulation system (both solute and solvent) that were reduced uniformly by tenfold ^33, 85^, and (x) default values of all other inputs of the PMEMD module. The FF12MC parameters are available in the Supporting Information of previous literature ^85^. The complex conformations were saved at 100-ps intervals in all simulations. The cluster analysis of all saved complex conformations was performed using the CPPTRAJ module of AmberTools 16 (University of California, San Francisco) with the average-linkage algorithm ^89^ (epsilon = 2.0 Å; RMS on all alpha carbon atoms of Residues 32–51, 60–97, and 108–124 of TREM2 and the entire LCD fragment). Centering the coordinates of the complex and imaging the coordinates to the primary unit cell were performed prior to the cluster analysis. The most populated conformation of the MD simulations identified by the cluster analysis was used as the simulation-refined model of the complex shown in Fig. 7.

### Surface Plasmon Resonance (SPR)

Affinity analysis was carried out using a Biacore T200 instrument (GE Healthcare Life Sciences) at Creative biolab. hTREM2 protein (Sino Biological, 11084-H08H) was directly immobilized on the CM5 chip using an amine coupling kit (GE Healthcare Life Sciences). Before immobilization, the CM5 sensor surface was activated using a mixture of 400 mM 1-ethyl-3-(3-dimethylaminopropyl) carbodiimide (EDC) and 100mM N-hydroxysuccinimide (NHS). Then, 10 μg/mL of hTREM2 in immobilization buffer (10 mM NaAc (pH 4.5)) was then injected into Fc2 sample channel at a flow rate of 10 μL/min. The amount of ligand immobilized was about 5,000 RU. The chip was deactivated by 1 M Ethanolamine hydrochloride-NaOH (GE Healthcare Life Sciences) at a flow rate of 10 μL/min for 420 s. The reference Fc1 channel underwent similar procedures but without injecting the ligand. The hTDP-43 protein (R&D, AP-190-100) was serially diluted with the running buffer to give a concentration of 1000, 500, 250, 125, 62.5, and 0 nM, respectively. Different concentrations of hTDP-43 proteins were then injected into Fc2-Fc1 channels at a flow rate of 30 μL/min, with a contact time of 120 s, followed by a dissociation time of 180 s. After each cycle of interaction analysis, the analyte injection, the association and dissociation process are all handling in the running buffer. The 4 TDP43 peptides (Mayo Clinic Rochester Proteomics Core) were serially diluted with the running buffer to give a concentration of 200, 100, 50, 25, 12.5, 6.25 and 0 μM, respectively. Different concentrations of TDP43 peptides were then injected into Fc2-Fc1 channels at a flow rate of 30 μL/min, with a contact time of 60 s, followed by a dissociation time of 90 s. After each cycle of interaction analysis, the analyte injection, the association and dissociation process are all handling in the running buffer. Data analysis was performed on the Biacore T200 computer and with the Biacore T200 evaluation software, using the 1:1 binding model.

### Statistics

Statistical details of the experiments, including sample sizes and statistical tests are described in figure legends. Results are displayed as mean ± SEM. All statistical analysis was performed with GraphPad Prism software (version 8). Samples sizes are comparable to similar studies ^25, 42, 87^. No statistical methods were used to predetermine sample sizes. All data were normally distributed and standard deviations were calculated to determine whether the data met assumptions of the statistical approach. Mean values for multiple groups was compared using a two-way ANOVA followed by Tukey’s *post hoc* test. Comparison of two groups was performed using a two-tailed unpaired Student’s *t*-test. P values < 0.05 were considered as statistically significant.

## Data availability

Further information and requests for resources and reagents should be directed to and will be fulfilled by the lead contact, Dr. Long-Jun Wu.

## Acknowledgments

The authors thank Dr. Marco Colonna (Washington University) for providing TREM2 KO mice; Dr. Ronald L. Klein (Louisiana State University) for providing AAV virus, Dr. Vanda A. Lennon (Mayo Clinic Rochester) for thoughtful discussion and manuscript editing; Dr. Wilfried Rossoll (Mayo Clinic Florida) for insightful suggestions; Dr. Aaron J. Johnson (Mayo Clinic Rochester), Dr. Yi Zhu (Mayo Clinic Rochester) and Dr. Kevin D. Pavelko (Mayo Clinic Rochester Immune Monitoring Core) for assistance with CyTOF sample preparation and data analysis; Dr. Akhilesh Pandey (Mayo Clinic Rochester Proteomics Core) for the discussion on LC-MS/MS experiment design and data analysis; Dr. Charles Howe and Dr. Benjamin Clarkson (Mayo Clinic Rochester) for providing iPSC-derived neurons; Dr. Richard M Weinshilboum (Mayo Clinic Rochester) for assistance with biochemistry experiments. The current study is supported by NIH grants R21AG064159 and R01NS088627 to L.J.W and R01AG066395 to G.B and N.Z.

## Author contributions

M.X., Y.U.L. and L.-J.W. designed the experiments and wrote the manuscript; M.X. and Y.U.L. performed most of the experiments; S.Z., Y.L., L.Z., U.S., Y.Z., N.Z., C.C.L performed some of the experiments; M.X. and D.B.B. performed data analysis; Y.-P.P. designed, performed, and analyzed the MD simulation study; J.Z performed and analyzed the mass spectrometry. Y.A.M., L.W., D.W.D., M.P.M. and G.B. provided resources for specific experiments.

## Extended Data Figure Legends

**Extended Data Fig 1.**
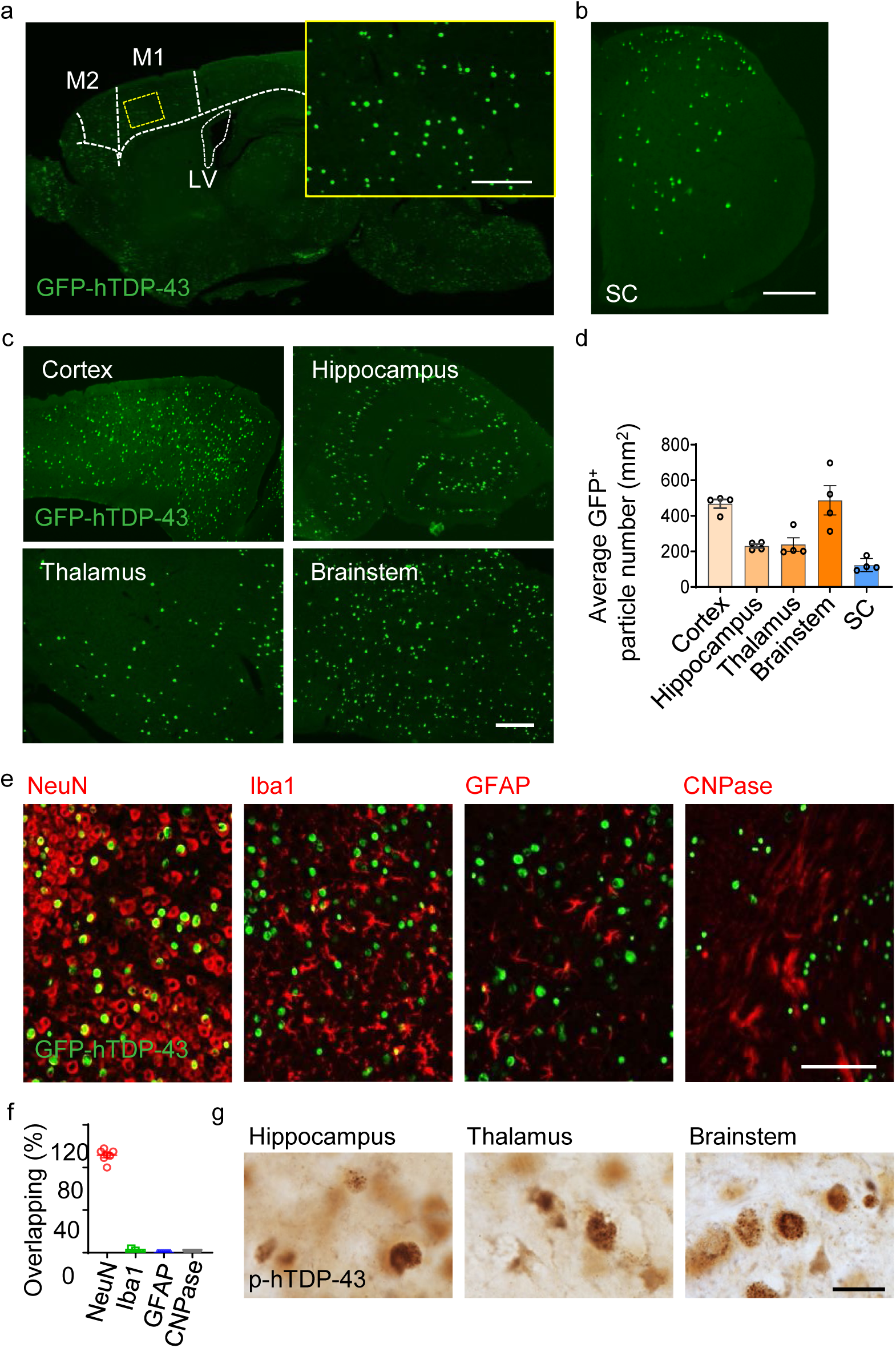
Characterizations of hTDP-43 expression in a neonatal TDP-43 mouse model. GFP-hTDP-43 expression was induced via intracerebroventricular injection of AAV9.CAG.hTDP-43-GFP in neonatal mice (AAV9.CAG.GFP as control). **a,** Representative image of brain GFP-hTDP-43 expression in C57/BL6 (WT) mice at 21 days post-infection (dpi). Primary motor cortex (M1), supplementary motor area (M2) and lateral ventricle (LV) are separated by dashed line. Inset shows seeding hTDP-43 expression at higher magnification as indicated by the area in dotted yellow box; Scale bar, 100 µm. **b,** Representative image of spinal cord GFP-hTDP-43 expression in C57/BL6 (WT) mice at 21 days post-infection (dpi); Scale bar, 200 µm. **c,** Representative image of brain GFP-hTDP-43 expression in the brain regions of cerebral cortex, hippocampus, thalamus and brainstem in WT mice at 21 days post-infection (dpi); Scale bar, 100 μm. **d,** Quantification of GFP-hTDP-43 expression in the brain regions of cerebral cortex, hippocampus, thalamus, brainstem and spinal cord (SC) in WT mice at 21 days post-infection (dpi). **e,** Representative images of co-immunostaining of GFP-hTDP-43 (green) with NeuN (red), Iba1 (red), GFAP (red) or CNPase (red) in the primary motor cortex of WT mice expressing hTDP-43 at 21 dpi. **f,** Quantification of the percentage of co-localization of GFP-hTDP-43 with NeuN, Iba1, GFAP or CNPase in the primary motor cortex of WT mice expressing hTDP-43 at 21 dpi (n = 7 per group). **g**, Representative images of p-hTDP-43 expression in the brain regions of hippocampus, thalamus, brainstem at 35 dpi; Scale bar, 20 µm. Data represented as mean ± SEM.

**Extended Data Fig 2.**
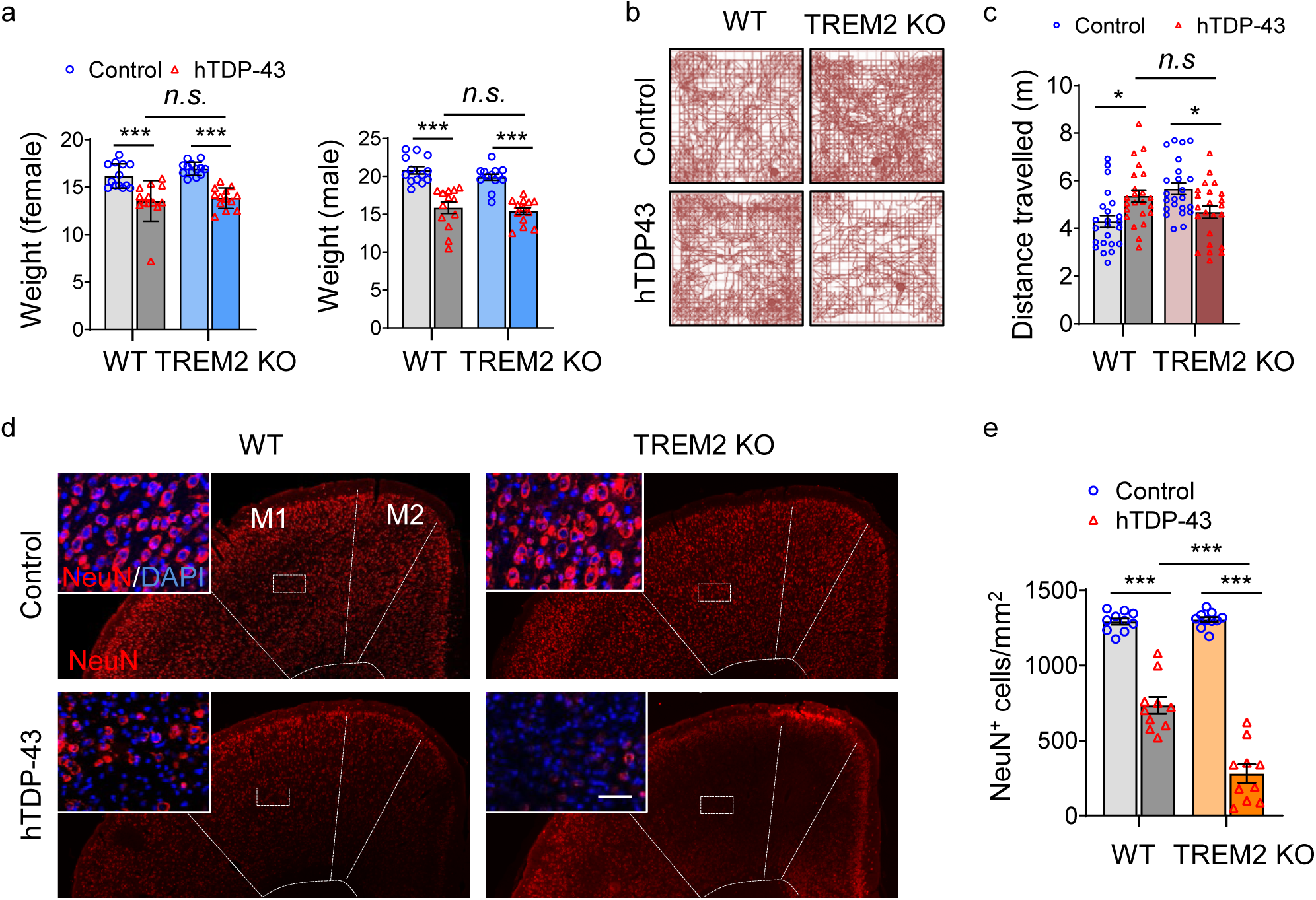
Characterizations of motor deficits and neuronal loss in a neonatal TDP-43 mouse model. GFP-hTDP-43 expression was induced via intracerebroventricular injection of AAV9.CAG.hTDP-43-GFP in neonatal mice (AAV9.CAG.GFP as control). **a,** Body weights were measured separately for male and female in indicated groups at 35 dpi (n = 13 per group). **b,** Representative traces of locomotor activity in an open field test at 35 dpi. **c,** Average total distance traveled during open field test (n=25 per group). **d,** Representative immunostaining images of NeuN (red) with DAPI (blue) in the primary motor cortex of indicated groups at 35 dpi. M1 and M2 are separated by dashed line. Insets show NeuN staining in layer 5 of primary motor cortex at higher magnification as indicated by the area in dotted white box; Scale bar, 20 µm. **e,** Quantification of NeuN positive cells in the primary motor cortex of indicated groups at 35 dpi (n=10 per group). Data represented as mean ± SEM. Significance was calculated using Two-way ANOVA, Tukey’s post-hoc analysis; *n.s.*, not significant; *** *P* < 0.001.

**Extended Data Fig 3.**
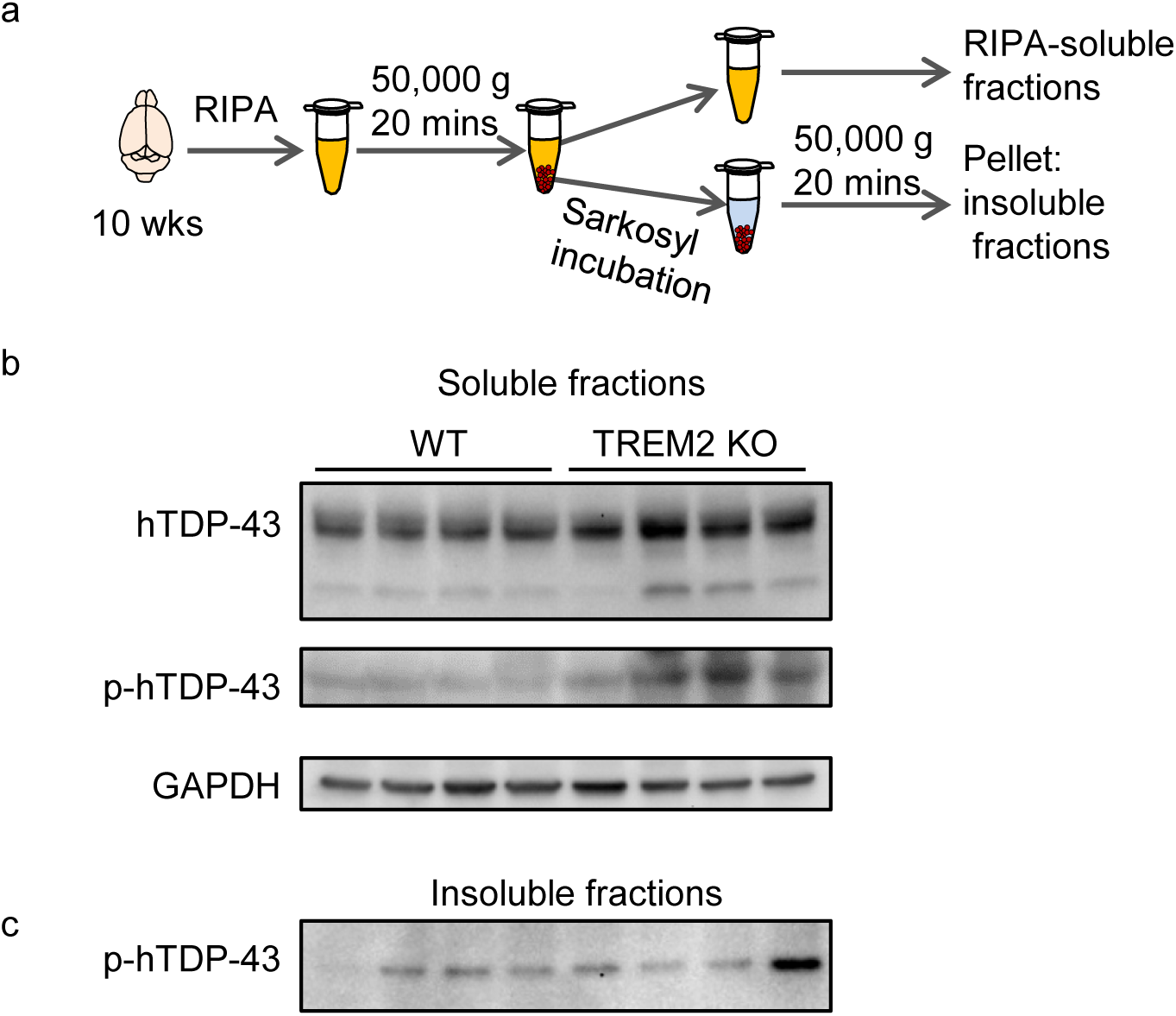
TREM2 deficiency leads to increased accumulation of hTDP-43 protein. hTDP-43 protein expression was induced via intracerebroventricular injection of AAV9.CAG.hTDP-43 in neonatal mouse (AAV9.CAG.Empty as control). **a,** Schematic representation of the velocity sedimentation protocol for separating soluble from Sarkosyl-insoluble aggregated proteins from an aqueous homogenate of brain tissue. **b,** Whole brain hTDP-43 and p-hTDP-43 (Ser403/404) immunoblots of the soluble fraction of WT and TREM2 KO mice at 70 dpi. **c,** Whole brain p-hTDP-43 immunoblots of the Sarkosyl-insoluble fraction of WT and TREM2 KO mice at 70 dpi. Western blots were independently repeated twice with n=4 for each group.

**Extended Data Fig 4.**
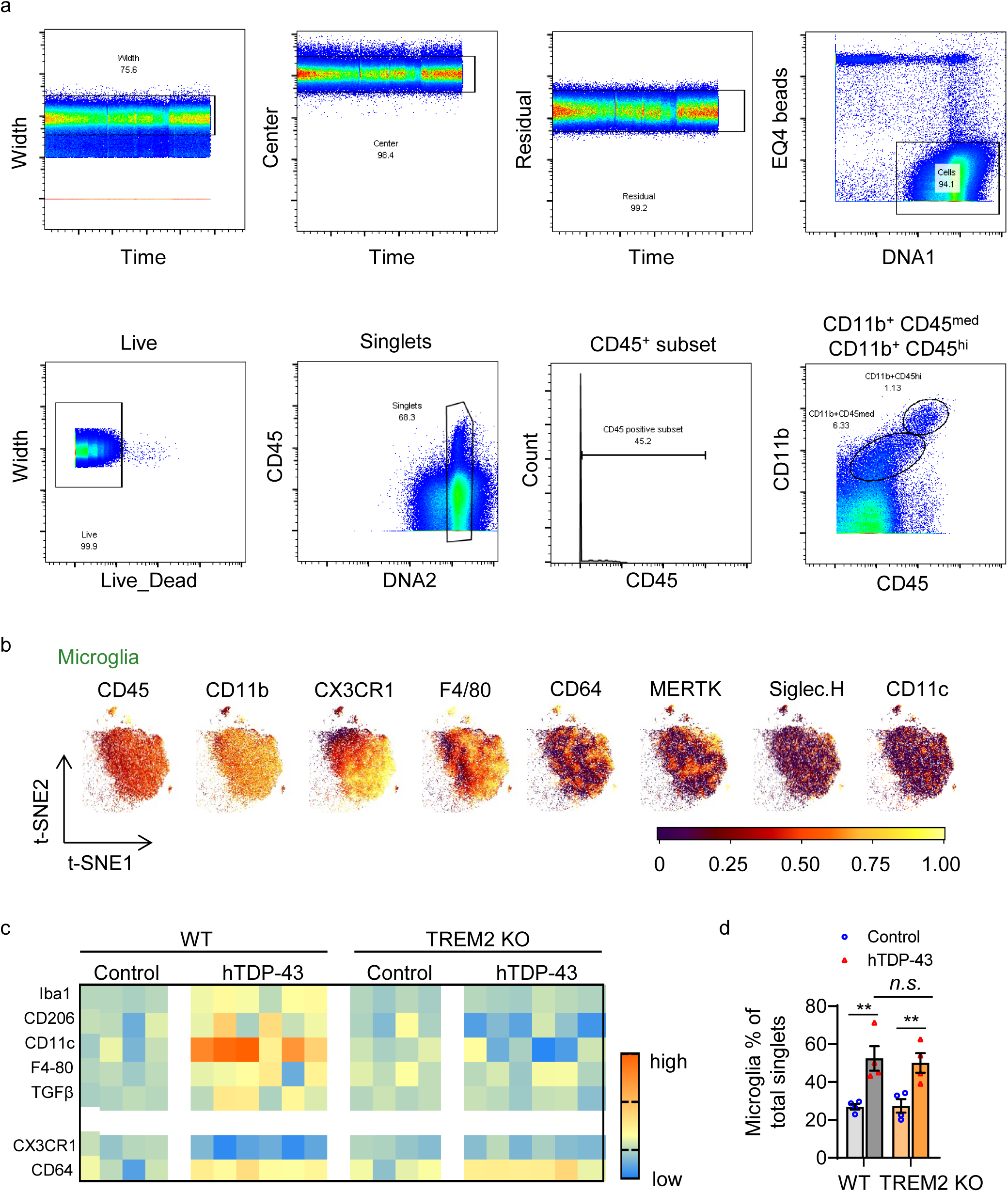
Phenotypic diversity of microglia in response to TDP-43 pathology by CyTOF. a, A representative gating strategy illustrating brain myeloid cell being subgated to CD45^med^CD11b^+^ microglia. **b,** Microglia were plotted onto a t-SNE. Plots represent distinguishing cell surface markers for microglia of 6-8 week-old WT mice. Clustering analysis revealed a major microglia population characterized by CD45^mid^:CD11b^+^:CX3CR1^hi^:F4/80^+^:CD64^+^:MERTK^+^:Siglec-H^+^:CD11c^-^. **c,** Heat map shows the change of expression levels of typical microglial markers in response to TDP-43 pathology in individual samples. Heat colors of expression levels have been scaled for each marker (blue, low expression; orange, high expression). **d,** Frequency analysis of microglia based on manual gating of indicated groups at 35 dpi (n=4 per group). Increased number of microglia was observed in both WT and TREM2 KO mice expressing hTDP-43.

**Extended Data Fig 5.**
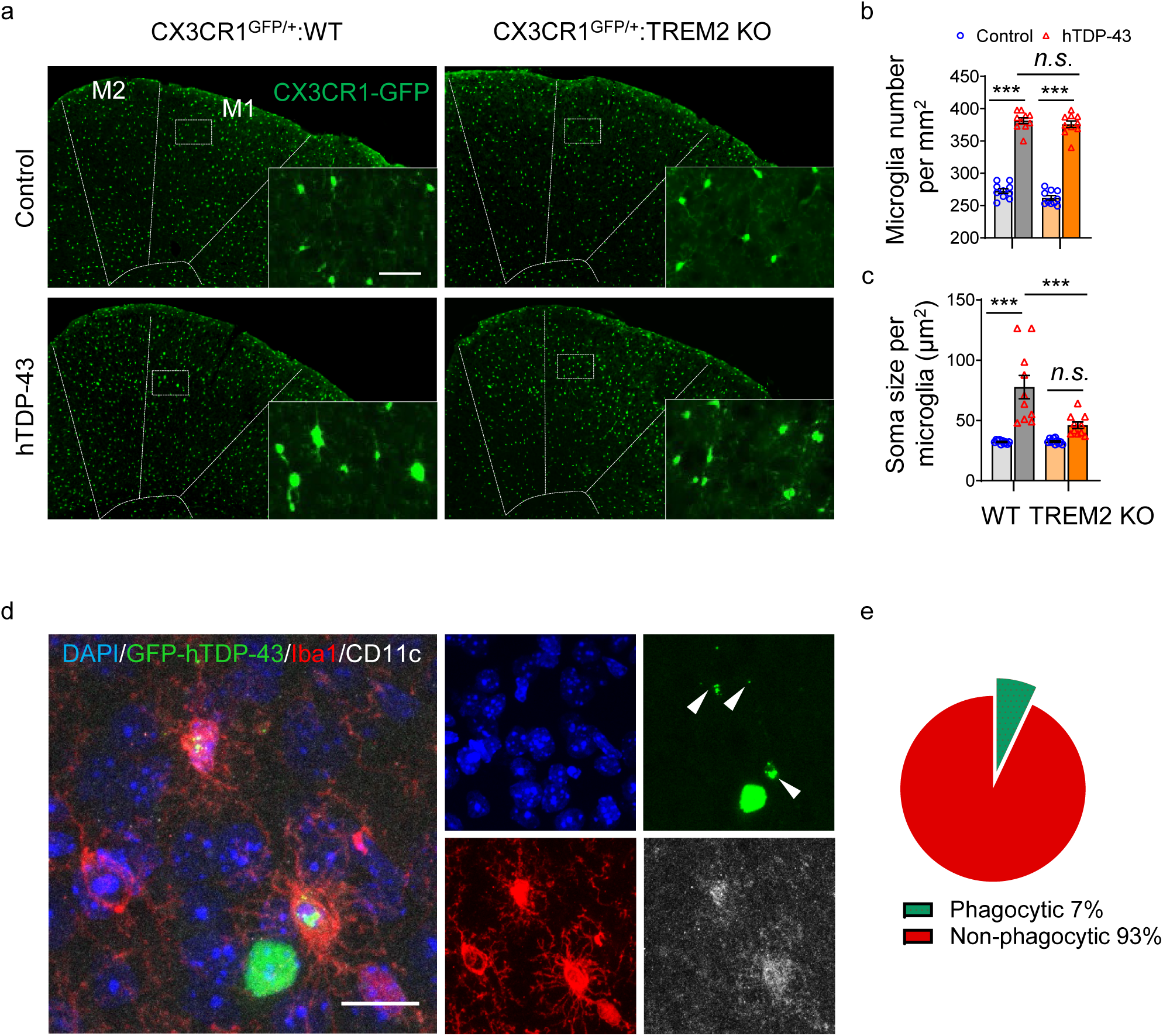
TREM2 deficiency attenuates hTDP-43-induced microglial activation. hTDP-43 protein expression was induced via intracerebroventricular injection of AAV9.CAG.hTDP-43 in neonatal mouse (AAV9.CAG.Empty as control). **a,** Representative images of GFP-expressing microglia in the primary motor cortex of indicated groups at 35 dpi. M1 and M2 are separated by dashed line. Insets show microglia at higher magnification as indicated by the area in white box; Scale bar, 50µm. **b, c,** Quantification of GFP-expressing microglia number (**b**) and soma size (**c**) in the primary motor cortex of indicated groups at 35 dpi (n = 10 per group). hTDP-43 increased microglia number in both CX3CR1^GFP/+^:WT and CX3CR1^GFP/+^:TREM2 KO mice. However, increased soma size was observed only in hTDP-43 CX3CR1^GFP/+^:WT mice. **d,** Representative images of co-localization of CD11c (white) with Iba1 (red) in microglia phagocytosing GFP-hTDP-43 (green) in the primary motor cortex of WT group at 35 dpi. White arrowheads indicate phagocytic puncta of GFP-hTDP-43; Scale bar, 20 µm. **e,** Pie chart representing the percentage of microglia phagocytosing GFP-hTDP-43 (green, phagocytic microglia) and those non-phagocytic microglia in the primary motor cortex of WT group at 35 dpi. Data represented as mean ± SEM. Significance was calculated using two-way ANOVA followed by Tukey’s post hoc test; *n.s.*, not significant; *** *P* < 0.001.

**Extended Data Fig 6.**
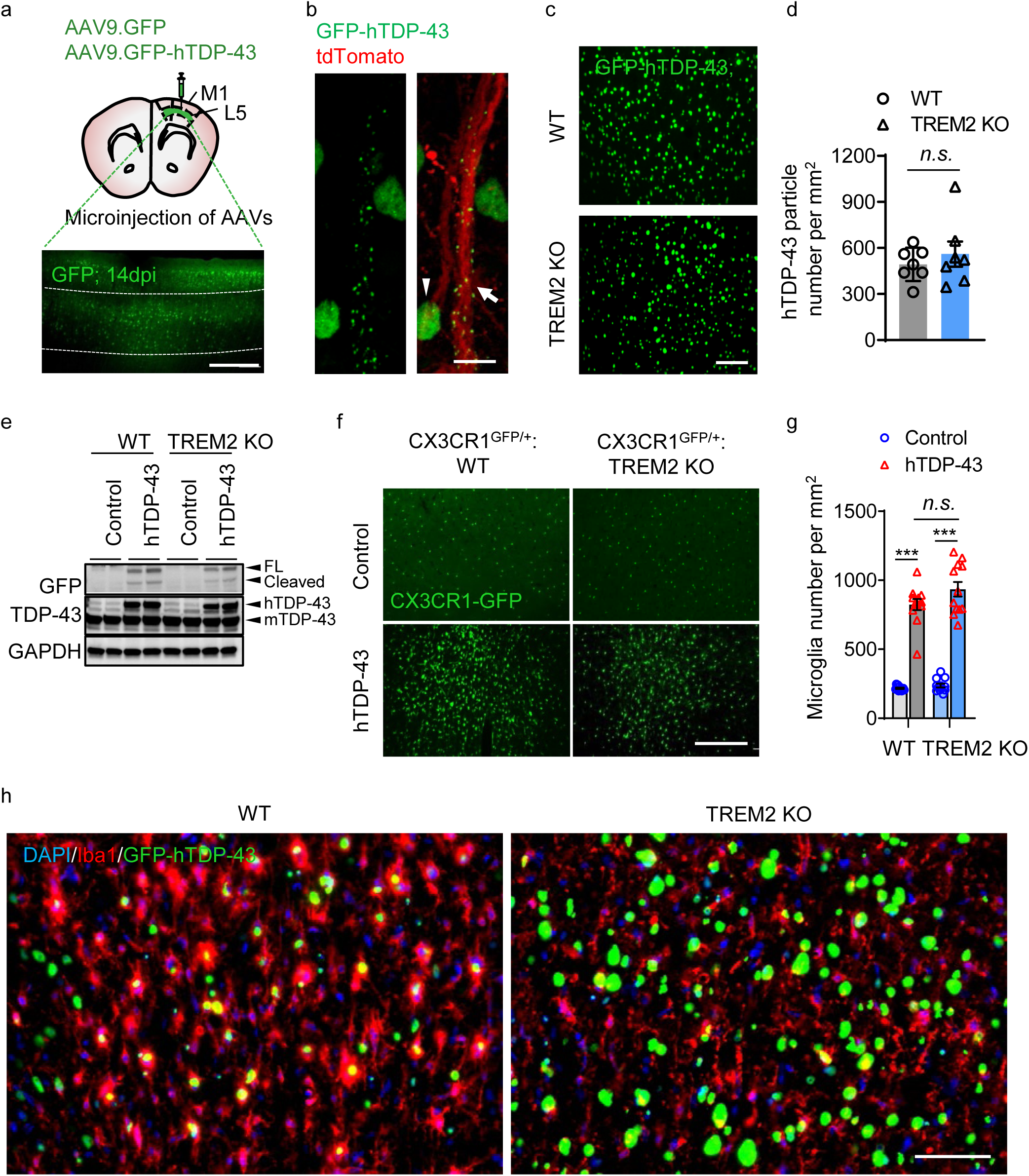
Characterizations of hTDP-43 expression mouse model via local virus injection in the primary motor cortex of adult mice. GFP-hTDP-43 or hTDP-43 was expressed in the primary motor cortex of 2-month-old mice via stereotactic intracerebral injection of AAV9.CAG.hTDP-43-GFP or AAV9.CAG.hTDP (AAV9.CAG.GFP or AAV9.CAG.Empty as control). **a,** Schematic picture showing stereotactic virus injection site (upper). Representative image of GFP expression in the primary motor cortex of 2-month-old WT mice at 14 dpi (lower); Dashed lines indicate the borders of layer 4&5; Scale bar, 100 µm. **b,** Representative images of translocation of GFP-hTDP-43 from nuclear to cytosol (white arrowhead) and dendrites (white arrow) in motor neurons of 2-month-old WT mice expressing GFP-hTDP-43 via stereotactic intracerebral injection at 14 dpi. AAV1.CAG.tdTomato virus was co-injected to visualize neurons; Scale bar, 5 µm. **c,** Representative images of GFP-hTDP-43 expression in the primary motor cortex at 14 dpi; Scale bar, 100µm. **d,** Quantification of GFP-hTDP-43 density in the primary motor cortex of WT and TREM2 mice at 14 dpi (n=7 per group). **e,** GFP-hTDP-43 immunoblots of primary motor cortex of WT and TREM KO mice at 14 dpi. Western blots were independently repeated four times (n=8 per group). GAPDH was used as loading control. **f,** Representative images of GFP-expressing microglia in the primary motor cortex of indicated groups at 14 dpi; Scale bar, 100µm. **g,** Quantification of GFP-expressing microglia number in the primary motor cortex of indicated groups at 14 dpi (n = 12 per group). **h,** Representative images of microglia (Iba1, red) phagocytosing GFP-hTDP-43 (green) in the primary motor cortex of WT but not TREM2 KO mice at 28 dpi; Scale bar, 100 µm. Data represented as mean ± SEM. Significances were calculated using student t test (**d**) and Two-way ANOVA, Tukey’s post-hoc analysis (**g**); *n.s.*, not significant; \**P* < 0.05, \*\**P* < 0.01, *** *P* < 0.001.

**Extended Data Fig 7.**
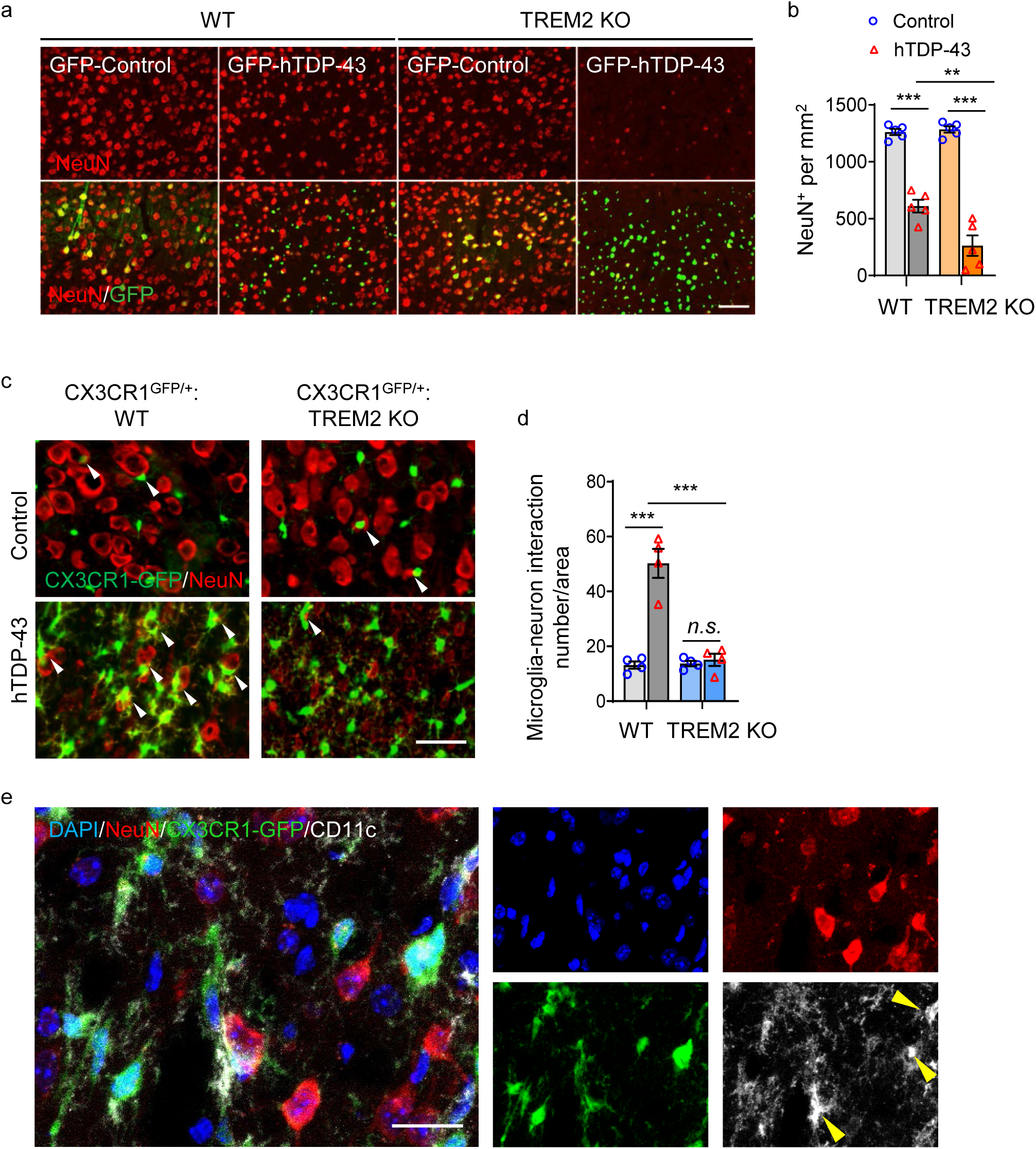
TREM2 deficiency facilitates hTDP-43-induced neurodegeneration. GFP-hTDP-43 or hTDP-43 was expressed in the primary motor cortex of 2-month-old mice via stereotactic intracerebral injection of AAV9.CAG.hTDP-43-GFP or AAV9.CAG.hTDP (AAV9.CAG.GFP or AAV9.CAG.Empty as control). **a,** Representative images of NeuN immunostaining (red) in the primary motor cortex of indicated groups at 28 dpi. Scale bar, 100 µm. **b,** Quantification of NeuN^+^ cells in the primary motor cortex of indicated groups at 28 dpi (n = 5 per group). hTDP-43 induced neurodegeneration was more prominent in TREM2 KO mice as compared with WT mice. **c,** Representative images of microglia (GFP) interaction with NeuN^+^ neurons (red) in the primary motor cortex of indicated groups at 28 dpi. Scale bar, 50 µm. **d,** Quantification of microglia-neuron interaction in the primary motor cortex of indicated groups at 28 dpi (n = 4 per group). Results show that hTDP-43 expression increased microglia and neuronal interaction in WT mice unlike in TREM2 KO mice. **e,** Representative images of CD11c (white) expression in microglia (CX3CR1-GFP, green) interacting with NeuN^+^ neurons (red) in the primary motor cortex of WT groups at 28 dpi. Scale bar, 20 µm. Data represented as mean ± SEM. Significance was calculated using two-way ANOVA followed by Tukey’s post hoc test; *n.s.*, not significant; \*\**P* < 0.01, *** *P* < 0.001.

**Extended Data Fig 8.**
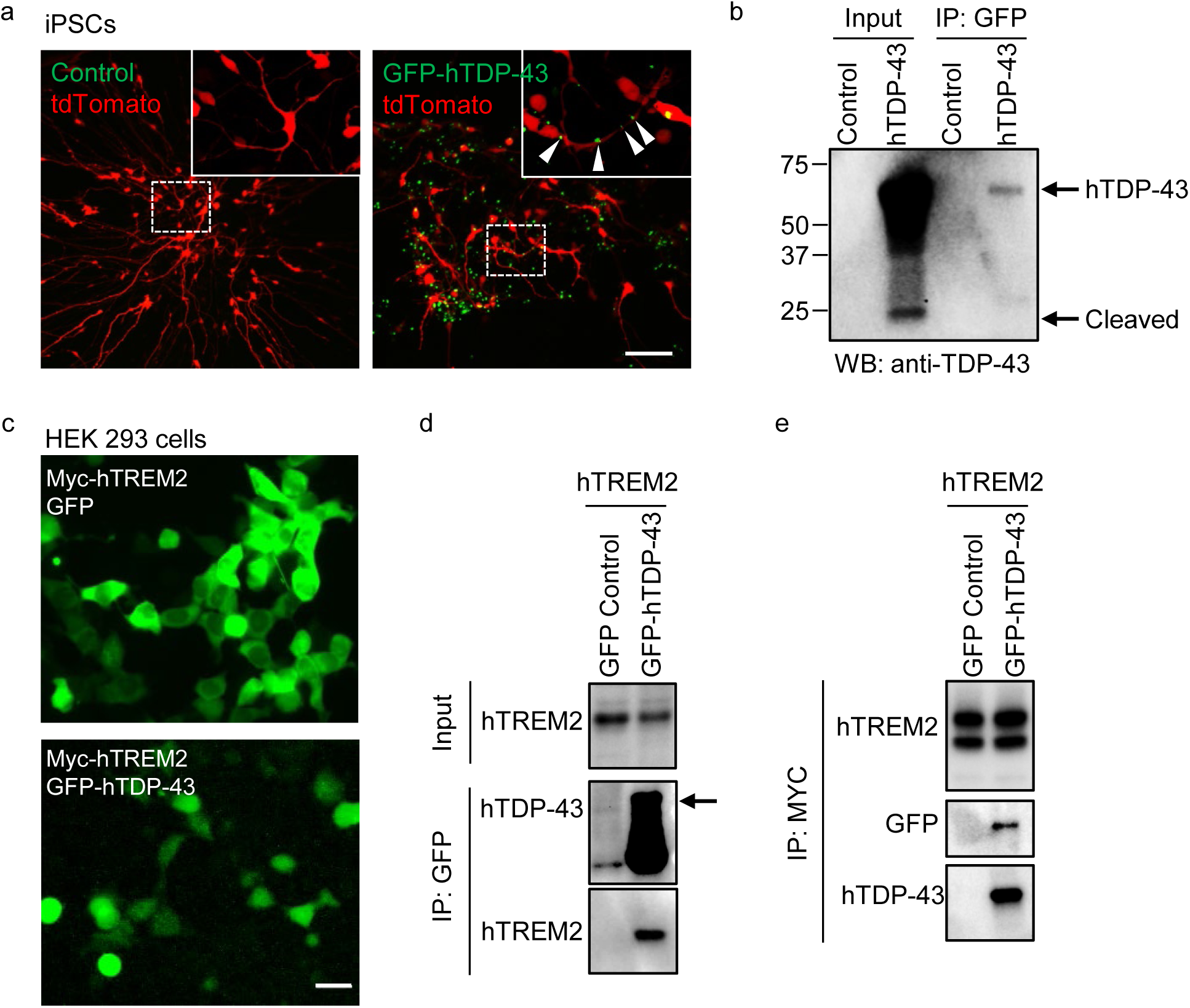
TDP-43 can be released from neurons and interact with TREM2 *in vitro*. a, Representative images of human iPSC derived neurons infected with AAV9.CAG.hTDP-43-GFP virus or control virus at 21dpi. AAV1.CAG.tdTomato virus used to visualize neurons; Scale bar, 100 µm. Inserts show neuron morphology at high magnification as indicated by the area in dotted white box. White arrowheads indicate GFP-hTDP-43 translocation in neurites. **b,** Immunoblots of TDP-43 contained within collected culture media of iPSCs (left) or in pulled down fractions by bead-immobilized GFP antibody (right). **c,** Representative images of HEK 293 cells co-transfected with myc-tagged human TREM2 (myc-hTREM2) and GFP-hTDP-43 C terminal fragment (residues 216-414) or GFP control plasmids. Scale bar, 20 µm. **d,** myc-hTREM2 was co-immunoprecipitated with GFP-hTDP-43 in HEK 293 cells using bead-immobilized GFP antibody. Experiments were independently repeated three times. **e,** GFP-hTDP-43 was co-immunoprecipitated with myc-hTREM2 in HEK 293 cells using bead-immobilized myc antibody. Experiments were independently repeated three times.

**Extended Data Fig 9.**
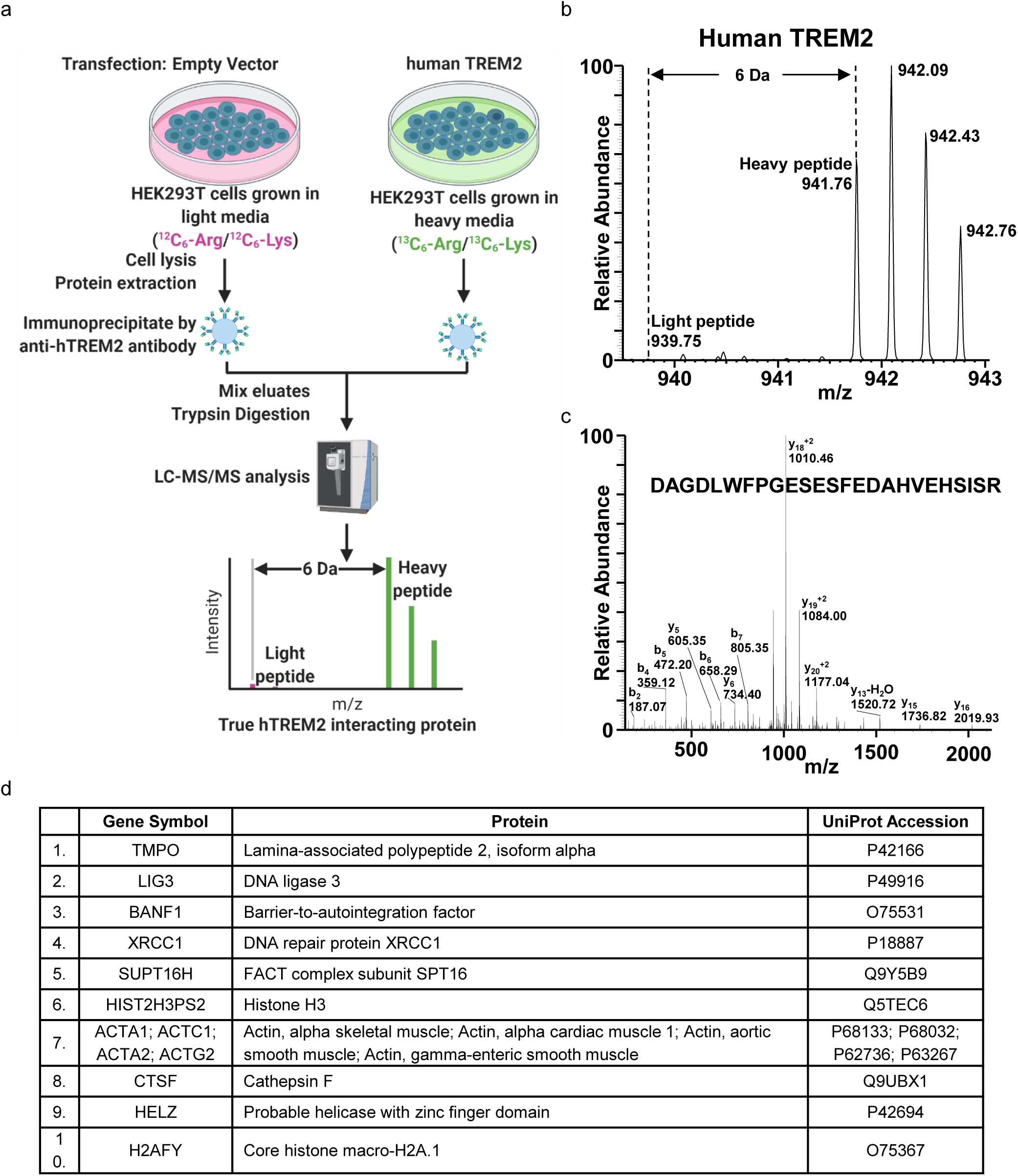
Human TREM2 interacting proteins in HEK293T cells. a, A schematic illustration of the SILAC methodology to identify human TREM2 interacting proteins in HEK293T cells (Created with BioRender.com). b and c, MS and MS/MS spectra of a peptide “D^137^AGDLWFPGESESFEDAHVEHSISR^161^” from human TREM2. d, A list of the 10 human TREM2 interacting proteins with highest intensity.

**Extended Data Fig 10.**
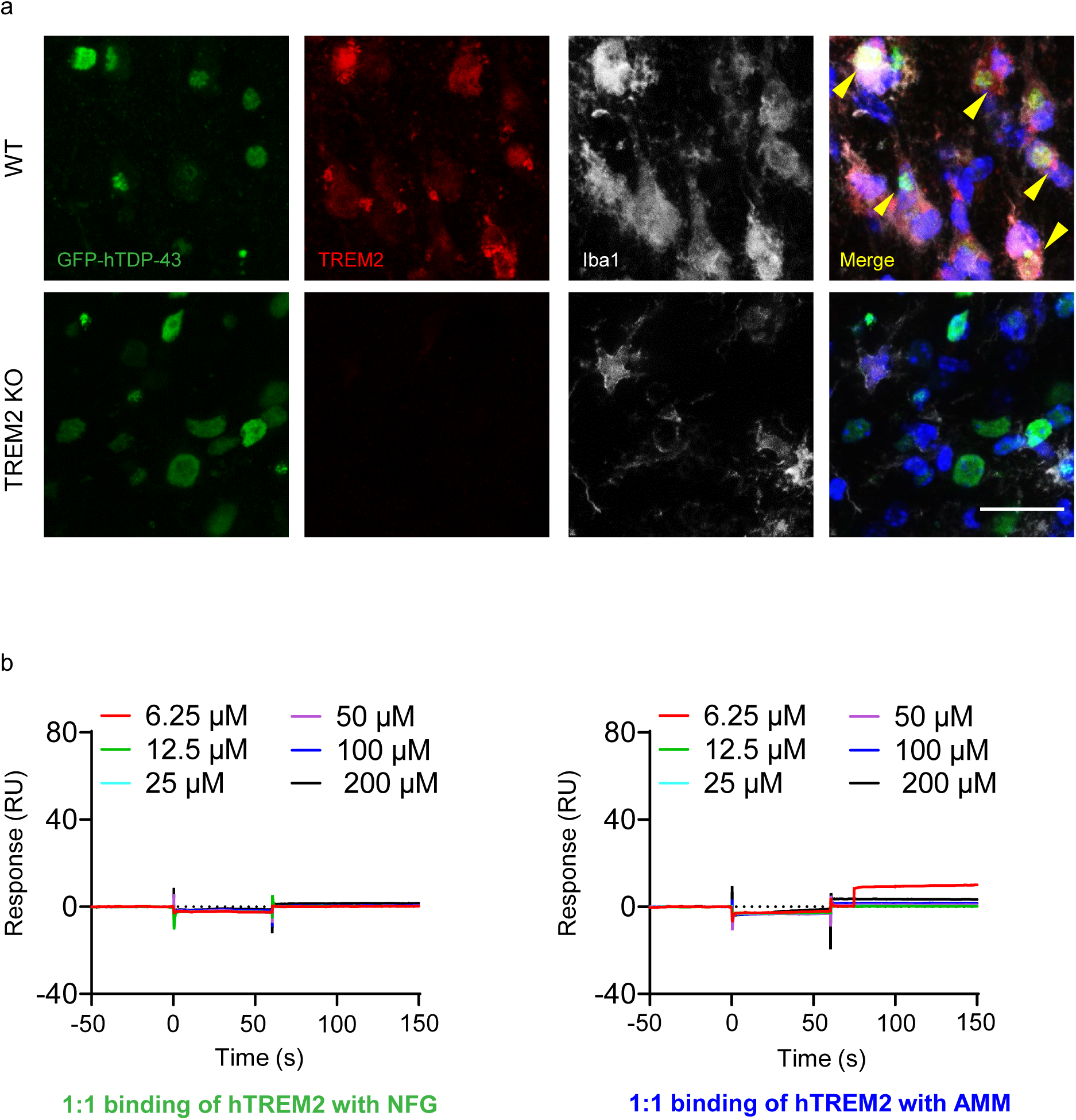
hTDP-43 interacts with TREM2 *in vivo* in mouse brain. GFP-hTDP-43 was expressed in the primary motor cortex of 2-month-old mice via stereotactic intracerebral injection of AAV9.CAG.hTDP-43-GFP. a, Representative images of co-localization of TREM2 (red) with Iba1 (white) in microglia phagocytosing GFP-hTDP-43 (green) in the primary motor cortex of indicated groups at 14 dpi. Arrowheads indicate co-localization of TREM2 with phagocytic puncta of GFP-hTDP-43; Scale bar, 20 µm. b, Recombinant hTREM2 ECD (Met1-Ser174) was immobilized on a CM5 BIAcore chip and binding of NFG and AMM at the indicated concentrations were determined by SPR.

**Extended Data Fig 11.**
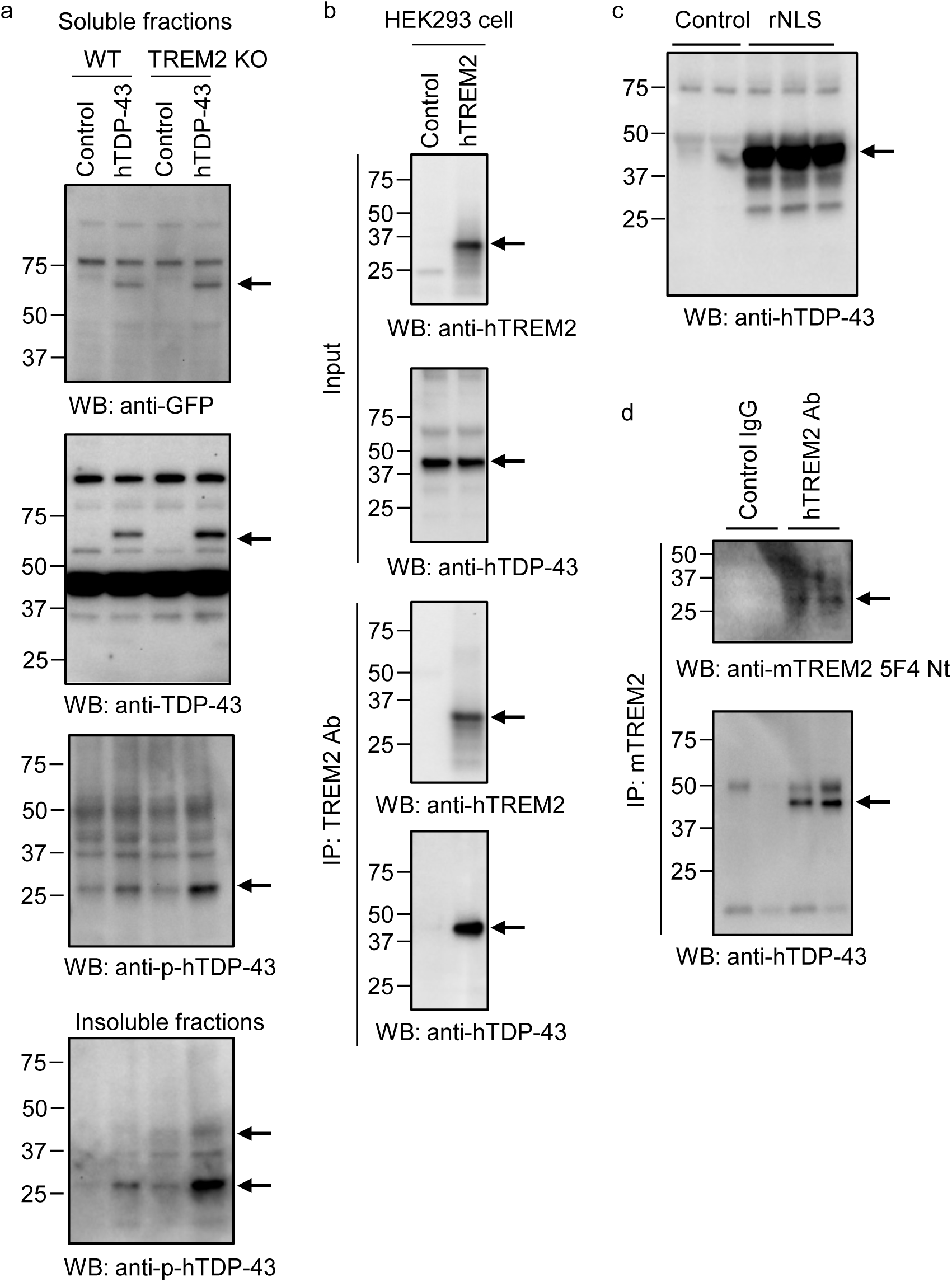
Full-length Western blot images. a, Full-length blot image for Figure 5g. **b,** Full-length blot image for Figure 6a. **c,** Full-length blot image for Figure 6f. **d,** Full-length blot image for Figure 6i.

**Extended Data Fig 12.**
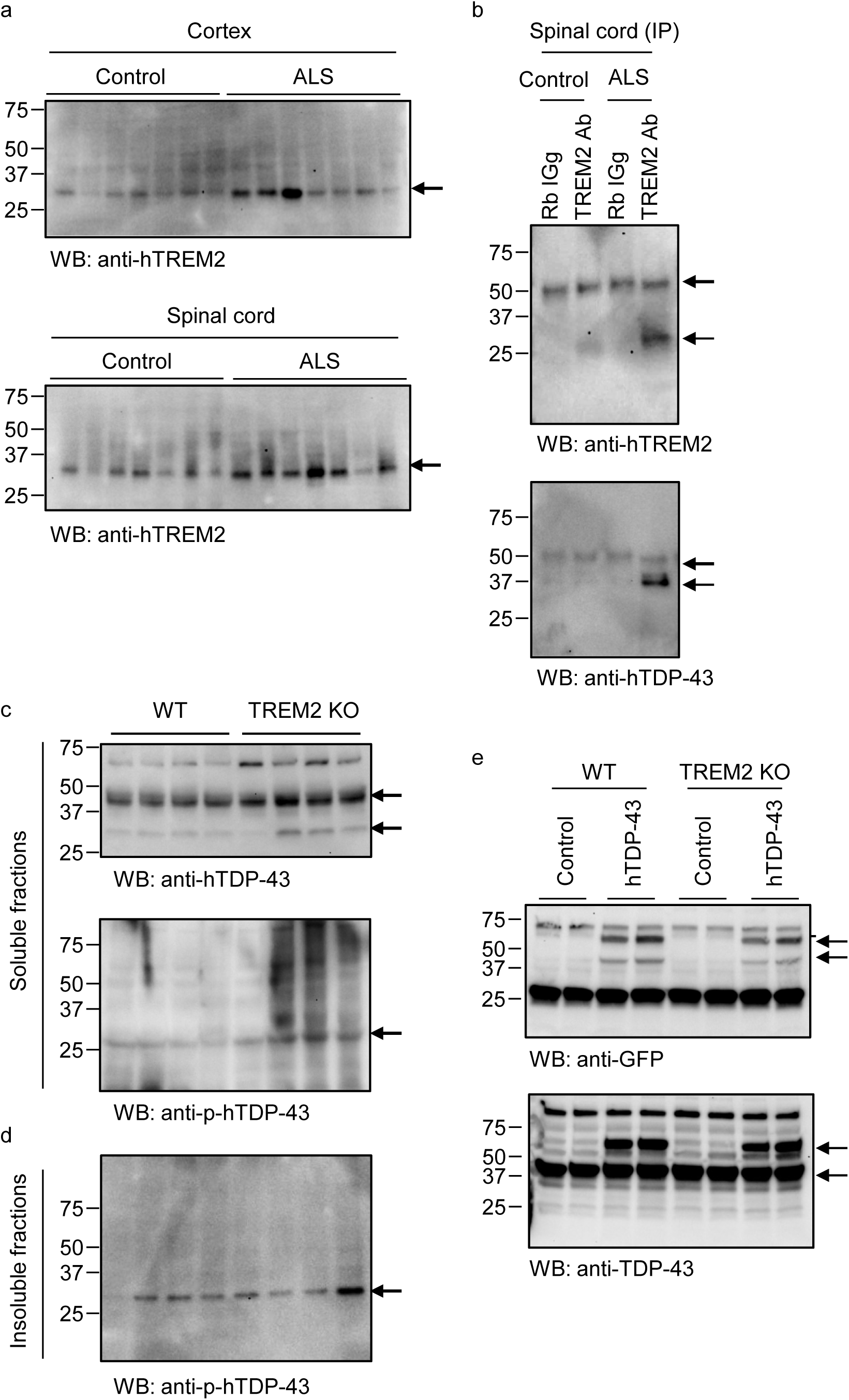
Full-length Western blot images. a, Full-length blot image for Figure 7a. **b,** Full-length blot image for Figure 7c. **c,** Full-length blot image for Extended Data Figure 3b. **d,** Full-length blot image for Extended Data Figure 3c. **e,** Full-length blot image for Extended Data Figure 6e.

**Extended Data Fig 13.**
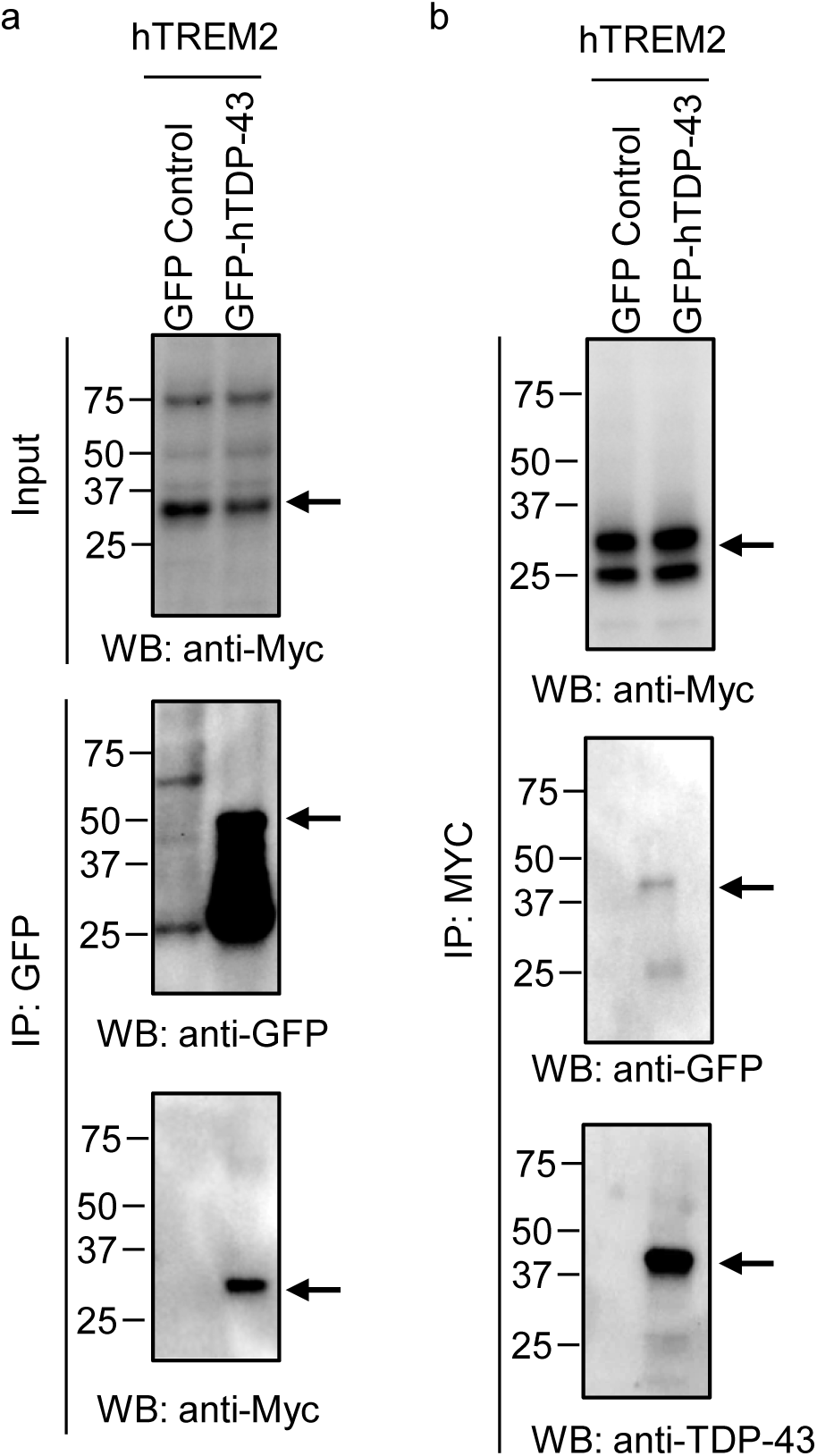
Full-length Western blot images. a, Full-length blot image for Extended Data Figure 8d. **b,** Full-length blot image for Extended Data Figure 8e.

